# Cell arrangement impacts metabolic activity and antibiotic tolerance in *Pseudomonas aeruginosa* biofilms

**DOI:** 10.1101/2023.06.20.545666

**Authors:** Hannah Dayton, Julie Kiss, Mian Wei, Shradha Chauhan, Emily LaMarre, William Cole Cornell, Chase J. Morgan, Anuradha Janakiraman, Wei Min, Raju Tomer, Alexa Price-Whelan, Jasmine A Nirody, Lars E.P. Dietrich

## Abstract

Cells must access resources to survive, and the anatomy of multicellular structures influences this access. In diverse multicellular eukaryotes, resources are provided by internal conduits that allow substances to travel more readily through tissue than they would via diffusion. Microbes growing in multicellular structures, called biofilms, are also affected by differential access to resources and we hypothesized that this is influenced by the physical arrangement of the cells. In this study, we examined the microanatomy of biofilms formed by the pathogenic bacterium *Pseudomonas aeruginosa* and discovered that clonal cells form striations that are packed lengthwise across most of a mature biofilm’s depth. We identified mutants, including those defective in pilus function and in O-antigen attachment, that show alterations to this lengthwise packing phenotype. Consistent with the notion that cellular arrangement affects access to resources within the biofilm, we found that while the wild type shows even distribution of tested substrates across depth, the mutants show accumulation of substrates at the biofilm boundaries. Furthermore, we found that altered cellular arrangement within biofilms affects the localization of metabolic activity, the survival of resident cells, and the susceptibility of subpopulations to antibiotic treatment. Our observations provide insight into cellular features that determine biofilm microanatomy, with consequences for physiological differentiation and drug sensitivity.

## INTRODUCTION

Chemical gradients form inevitably during multicellular growth, leading to the establishment of distinct subzones–with different conditions of resource availability–in cellular aggregates and complex organisms [1]. The conditions of these subzones are important because they promote physiological differentiation, in turn affecting morphogenesis, metabolic activity, and susceptibility to drugs [2–6]. The capacity for substrate uptake and transport are therefore critical features of a cellular aggregate that are linked to structural development and our ability to treat disease, with diverse mechanisms controlling these processes in various multicellular structures and organisms. Many macroscopic eukaryotes, for example, contain circulatory systems in which specialized cells form conduits that deliver water and other resources to cells [7]. These specialized transport systems allow for efficient nutrient allocation, enabling multicellular organisms to grow, develop, and maintain homeostasis.

The predominant multicellular structure of the microbial world is the biofilm, an aggregate of cells attached to each other and encased in a self-produced matrix [8]. There have been many studies that have uncovered the presence and importance of spatial organization within the biofilm [9–17]. Biofilms often form at interfaces and are subject to steep external and internal chemical gradients, raising questions about how nutrients are allocated efficiently within these structures. While evidence suggests that passive diffusion through the matrix is a significant mechanism of resource distribution within biofilms [18,19] observations of biofilms formed by model microbes also indicate that morphological features and differentiation could impact substrate transport [20–23].

In this study, we examined the cellular-level anatomy of macrocolonies formed by the bacterium *Pseudomonas aeruginosa*, which is a major cause of biofilm-based and chronic infections in immunocompromised individuals [24]. We found that *P. aeruginosa* biofilms contain vertically arranged zones that differ with respect to cellular orientation and the proximity of related cells. Genetic analyses and microscopic imaging allowed us to identify cell surface components required for this patterning. Importantly, these approaches also revealed that cellular orientation impacts substrate uptake and distribution across the biofilm. Additional analysis via stimulated Raman scattering (SRS) microscopy and an assay for cell death uncovered correlations between cellular-arrangement subzones, physiological status, and susceptibility to antibiotic treatment. Collectively, these data link microanatomy to metabolic differentiation--and its consequences for survival and drug resilience—within maturing biofilms.

## RESULTS AND DISCUSSION

### Mature *P. aeruginosa* biofilms contain vertical, clonal striations

Growth at surface-air interfaces has been used for decades to guide treatment decisions made in the clinical setting and is a focus of research into *P. aeruginosa* biofilm development [25–34]. In particular, our group and others have used macrocolony biofilms as models to study the molecular links between resource availability and multicellular physiology [35–38]; we have discovered a variety of pathways and mechanisms that support the biofilm lifestyle, some of which contribute to pathogenicity and antibiotic tolerance [39,40]. Macrocolony biofilms are formed when 1-10 uL of a cell suspension are pipetted onto agar-solidified media and incubated for several days (**Figure 1A**). The macrocolony biofilm model provides a high-throughput, macroscopic readout for the effects of a broad variety of factors–such as polysaccharide production, pilus function, and second messenger levels–that affect *P. aeruginosa* biofilm development [41–43]. It also provides a reproducible system for growing a physiologically differentiated multicellular structure that is amenable to techniques for mapping variations in extra- and intracellular chemistry, gene expression, and metabolic activity in situ [40,44–47].

**Figure 1.**
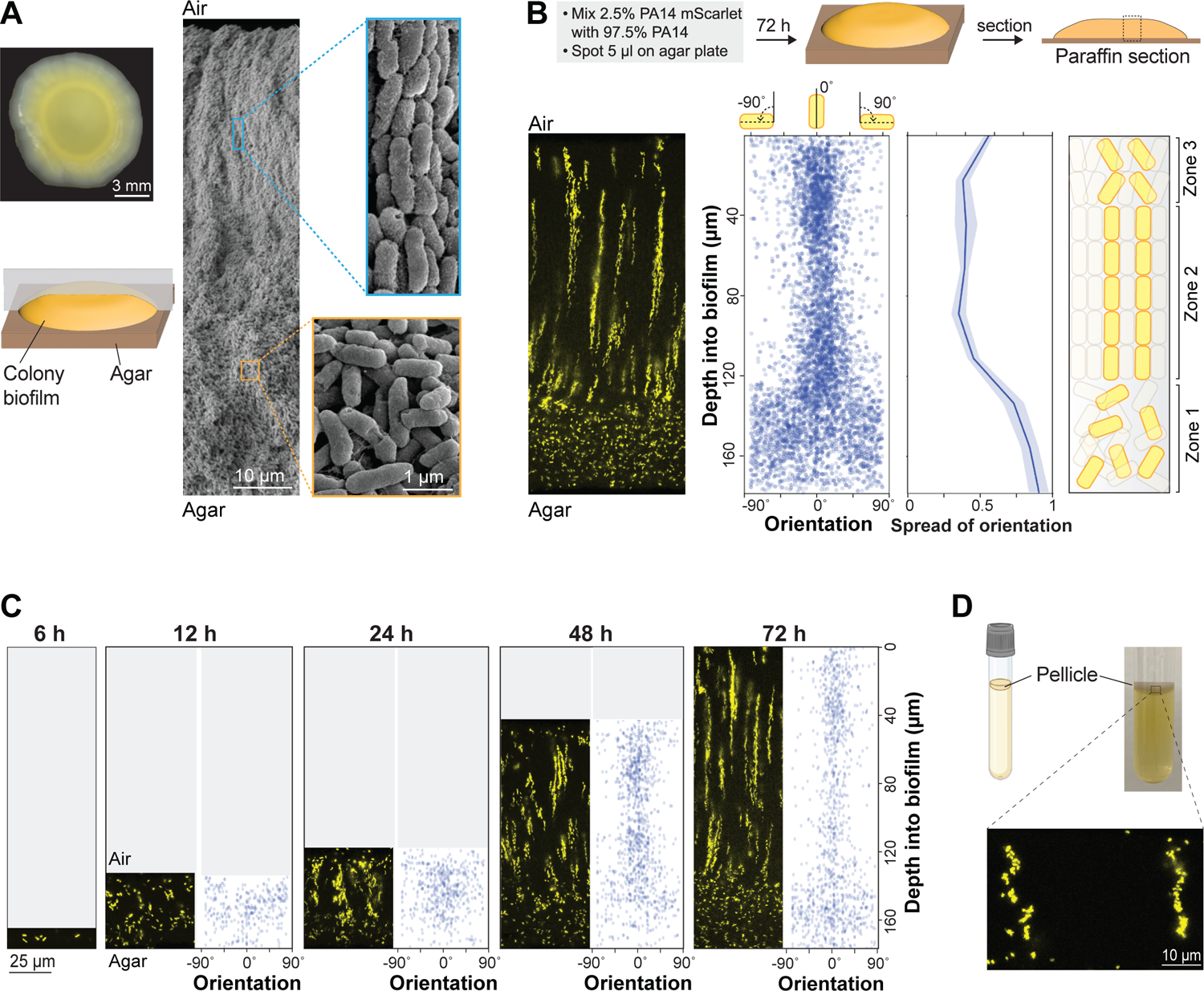
*P. aeruginosa* cells form vertical striations across depth in colony and pellicle biofilms. (**A**) **Left**: Top view of a *P. aeruginosa* colony biofilm grown for three days on 1% tryptone + 1% agar, and schematic showing orientation of the sample used for SEM imaging. **Right**: SEM images of a full colony biofilm cross-section. Insets of higher magnification show cellular arrangement and morphology for the indicated locations in the biofilm. (**B**) **Top**: Schematic of mixing assay method. **Left**: Fluorescence micrograph of a thin section prepared from a colony biofilm grown in the mixing assay. **Center**: Orientation across depth for fluorescent cells detected in biofilm thin section micrographs. The “spread of orientation” is the standard deviation of orientation values for each pixel across biofilm depth; the values shown in the plot are the average “spread of orientation” at each depth for thin section images taken from six biological-replicate biofilms. Shading represents the standard deviation for this average. **Right**: Schematic of cellular arrangement across depth in mature biofilms. (**C**) Micrographs of biofilms prepared as described in (B), but sacrificed at the indicated time points. (**D**) **Top**: Setup used to grow pellicle biofilms for microscopy. **Bottom**: Fluorescence micrograph of a thin section prepared from a pellicle biofilm. The inoculum contained 2.5% cells that constitutively express mScarlet. Images shown in this figure are representative of at least two independent experiments. mScarlet fluorescence is colored yellow.

Despite the significance of the macrocolony model to our understanding of *P. aeruginosa* biology, we lack detailed information about how cells are arranged within these structures and the genetic determinants of this arrangement. To begin to address this gap, we used scanning electron microscopy (SEM) to examine a WT *P. aeruginosa* PA14 macrocolony biofilm that had grown for 3 days on 1% tryptone, 1% agar. SEM images revealed two distinct zones, containing either “disordered” cells (in various orientations) or “ordered” cells (aligned lengthwise) (**Figure 1A**). Similar arrangements of cells aligned lengthwise with ordered packing have been described for diverse bacterial systems [11,48–56].

To assess the distribution of clonal (related) cells in the ordered region, we grew macrocolony biofilms in a “mixing assay” in which the inoculum contained 2.5% of a strain that constitutively expresses mScarlet as a fluorescent marker (P_PA1/04/03_-*mScarlet*). (The remaining 97.5% of the cell population was unmarked.) Biofilms were embedded in paraffin and thin-sectioned, and sections taken from the colony center (**Figure 1A**) were imaged by fluorescence microscopy [57] (**Figure 1B**). We discovered that fluorescent cells within the disordered region (zone 1) are randomly dispersed, while fluorescent cells in the ordered region are arranged in evenly spaced vertical striations reminiscent of the cellular packing seen in the SEM images (**Figure 1A**). To determine their orientations, we used a custom Python script that allowed us to detect individual cells and plot their angles relative to the vertical axis across biofilm depth. While the fluorescent cells in zone 1 are randomly oriented, those in zone 2 are oriented along the vertical axis, consistent with the arrangements visible by SEM (**Figure 1A**). Finally, in the uppermost portion of the biofilm (zone 3), cell orientation shows increased deviation from the vertical alignment, though fluorescent cells still appear to be spatially aggregated (**Figure 1B**).

Growth at the liquid-air interface constitutes another popular model system that has provided insights into mechanisms of biofilm formation [30,33,58,59]. To examine whether striated arrangement, similar to that observed in the macrocolony biofilm, develops in a biofilm grown at the liquid-air interface (pellicle) we used the same ratio of marked and unmarked cells as inocula for pellicle biofilm cultures. After three days of growth, pellicle biofilms were transferred to agar-solidified media and immediately embedded in paraffin and thin-sectioned. Microscopic analysis revealed that mixing-assay pellicle biofilms also show fluorescent striations (**Figure 1D**). Each of the pellicles imaged showed striations covering most or all of the biofilm depth and containing cells in random orientations. Together, these findings demonstrate that clonal subpopulations of *P. aeruginosa* cells arrange in vertical striations relative to the gel- or liquid-air interface during macrocolony or pellicle biofilm growth.

### Biofilm striations resolve and become longer over time

The presence of clonal striations in 3-day-old biofilms raised the question of how these features develop over the course of the incubation. To examine this, we grew replicate mixing-assay biofilms and sacrificed the biofilms for microscopic analysis at a series of time points.

Fluorescence imaging showed that PA14 cells in macrocolony biofilms are randomly oriented and distributed at 6 and 12 hours of incubation (**Figure 1C**). At 24 hours of incubation the upper two-thirds of the biofilm begin to show clustering and increased vertical orientation of fluorescent cells that gradually becomes more resolved over time. Between 24 and 72 hours of incubation, both zone 1 and zone 2 showed gradual increases in height (3-fold and 2.6-fold, respectively). These observations suggest that cells in zone 1 are able to move away from each other, while in contrast cells in zone 2 remain positioned end-to-end, after division (**Figure 1C**).

### Resource availability affects the organization of cellular-arrangement zones and profiles of metabolic activity in biofilms

Biofilms growing at interfaces form resource gradients as they get thicker due to diffusion limitation and consumption by cells closer to the boundary [1,60]. These gradients define chemical microniches that promote physiological differentiation. We hypothesized that changes in resource availability would alter cellular arrangement in *P. aeruginosa* macrocolonies. We tested this assumption by growing mixing-assay biofilms on different concentrations of tryptone and imaging thin-sections by fluorescence microscopy. As expected, biofilms grew thicker when we provided increasing amounts of tryptone, indicating that a vertical tryptone gradient formed across biofilm depth with limiting concentrations at the air interface (**Figure 2B**). We were surprised to find that on 0.25% tryptone fluorescent striations and vertically oriented cells were visible in the zone close to the biofilm base, while dispersed and randomly oriented cells were visible in the upper portion of the biofilm. In contrast, biofilms grown on 0.5% tryptone showed an overall zone organization similar to that seen on 1% tryptone. The ordered region of these biofilms, however, appeared more similar to zone 3 of 1%-tryptone grown biofilms in that cellular orientation showed more deviation from vertical and striations were thicker and not as well-segregated as in biofilms grown on the higher concentration (**Figure 2B**). This further suggests that the level of dispersion displayed by cells in a given zone does not correlate with the absolute concentration of tryptone: for example, the disordered zone corresponded to a tryptone-limited region in the 0.25%-tryptone-grown biofilm, while the disordered zones instead corresponded to the most tryptone-replete regions in the 0.5%- and 1%-tryptone-grown biofilms. These observations show that striation formation is sensitive to nutrient availability but suggest that the absolute concentration of tryptone is not the sole factor determining the organization of cellular-arrangement zones in biofilms.

**Figure 2.**
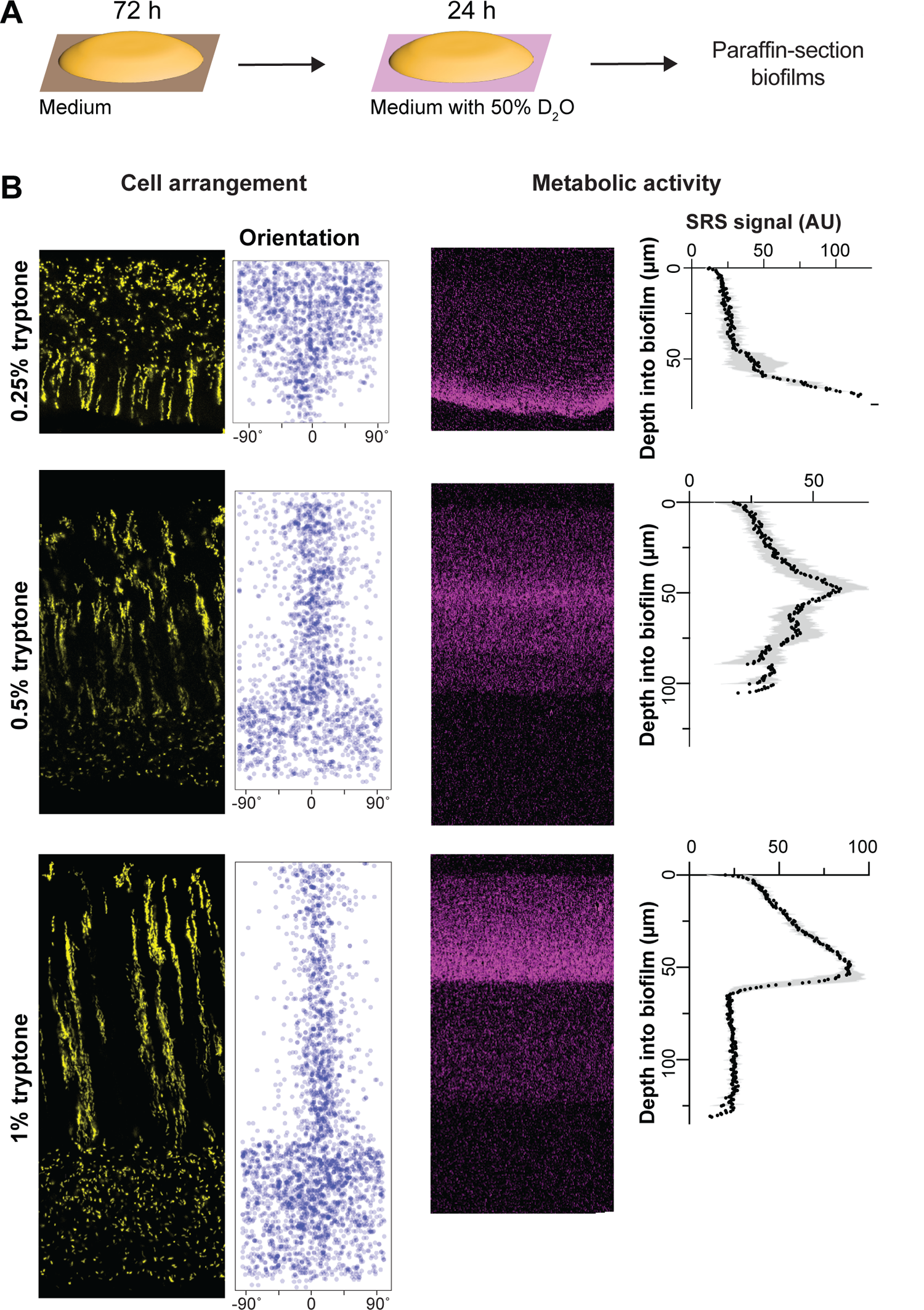
Resource availability affects the organization of cellular-arrangement zones and metabolic activity in biofilms. (**A**) Experimental setup for growing *P. aeruginosa* biofilms on agar plates and their subsequent transfer to medium containing D_2_O for analysis of metabolic activity by stimulated Raman scattering (SRS) microscopy. (**B**) **Left**: Fluorescence micrographs and quantification of cellular orientation across depth for thin sections from mixing-assay biofilms grown on different concentrations of tryptone. **Right**: SRS microscopy images and SRS signal across depth for thin sections from mixing-assay biofilms grown on different concentrations of tryptone. SRS signal represents the average of three biological replicates with shading indicating the standard deviation.

Because gradients within biofilms affect physiological status, we next sought to test whether the effect of tryptone concentration on cellular-arrangement zone properties and organization correlated with effects on metabolic activity. Our groups have established a protocol to visualize metabolic activity in intact biofilm thin sections using stimulated Raman scattering (SRS) microscopy [40]. In this method, biofilms are grown on agar plates, then transferred to a medium containing D_2_O for 24 hours (**Figure 2A**) before the preparation of thin sections. Incorporation of D into D-C bonds is an indicator of metabolic activity and is detected by SRS microscopy. We found that, for mixing-assay biofilms grown on each concentration of tryptone, metabolic activity peaked at a depth of 40-70 µm (**Figure 2B**). Microelectrode measurements have shown that *P. aeruginosa* biofilms grown at gel-air interfaces contain oxygen gradients [45]. The observation that maximal metabolic activity occurs at a consistent depth in biofilms grown at different tryptone concentrations suggests that maximum metabolic activity occurs where the relative concentrations of oxygen and tryptone are optimized [61]. Strikingly, the region with maximum metabolic activity overlapped with a region containing vertically oriented cells in biofilms grown at all tryptone concentrations (**Figure 2B**).

### Global regulators and cell surface components affect cell patterning

*P. aeruginosa* colony biofilm structure at the macroscale is determined primarily by the regulation of exopolysaccharide production in response to environmental cues and cellular signals [32,33,62–64]. To uncover the genetic determinants of biofilm cellular arrangement at the microscale, we conducted a targeted screen of 48 mutants lacking regulatory proteins, cell surface components, and other factors that contribute to WT biofilm formation and physiology in macrocolonies and other biofilm models (**Figure 3A**) [63–66]. For each of these mutants, a corresponding, constitutively fluorescent strain was generated that expressed the P_A1/04/03_-*mScarlet* construct in the mutant background. Mixed biofilms of each mutant were grown for three days and thin sections were prepared for microscopic analysis.

**Figure 3.**
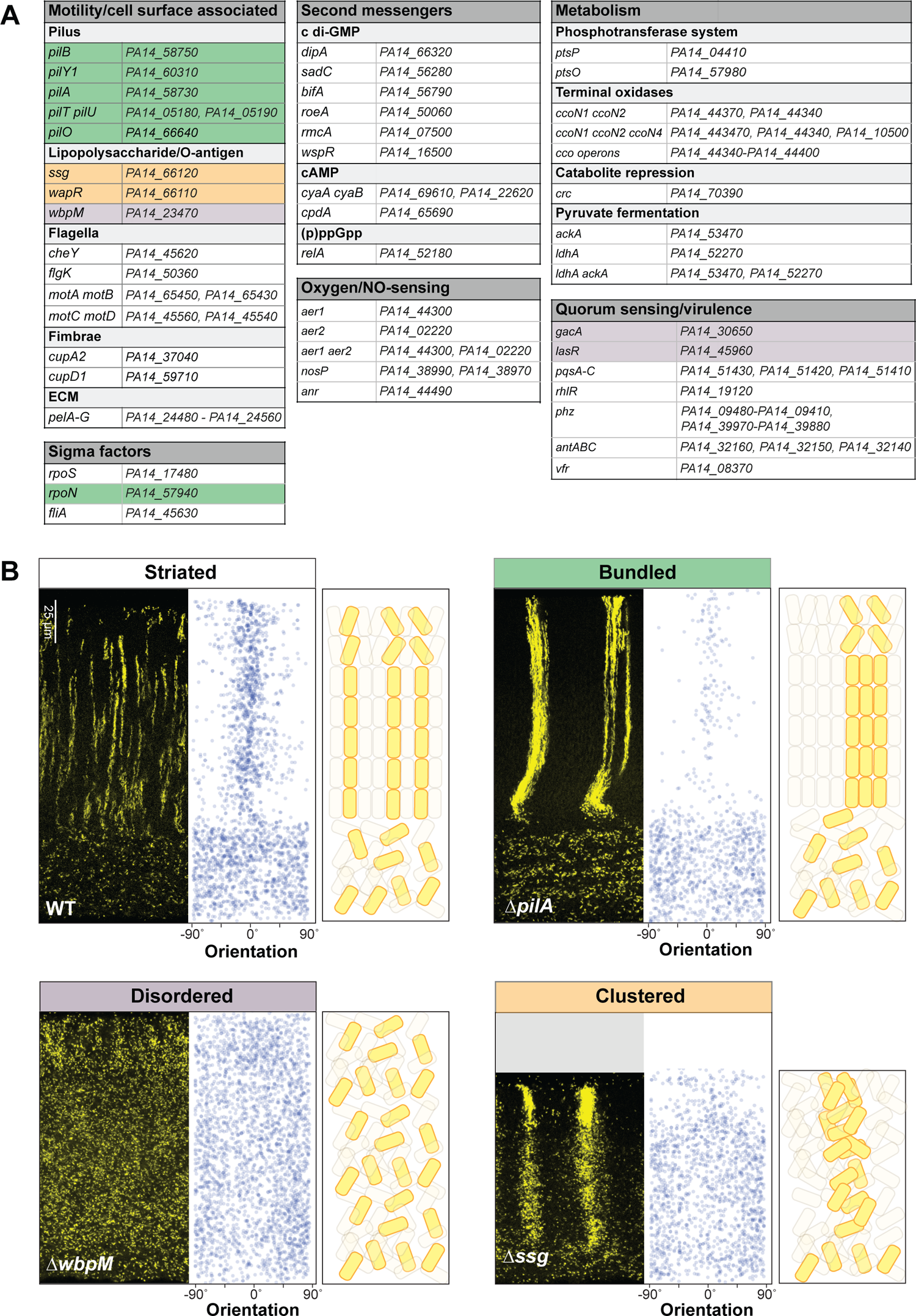
Specific global regulators and cell surface components are required for wild-type cell patterning in colony biofilms. (**A**) List of mutants screened for altered cellular arrangement across depth in biofilms. Those showing altered cellular arrangement are shaded and colors correspond to the phenotype categories shown in (B). (**B**) Fluorescence micrographs of thin sections from WT and indicated mutant biofilms grown on 1% tryptone + 1% agar for 3 days. Biofilm inocula contained 2.5% cells that constitutively express mScarlet. mScarlet fluorescence is colored yellow. Quantification of orientation across depth is shown for each image, and cartoons of cellular arrangement are shown for each phenotype category. Images shown are representative of at least two independent experiments.

Imaging allowed us to identify four classes of cellular arrangement phenotypes, which we refer to as “striated”, “bundled”, “disordered”, or “clustered” (**Figure 3B**). Thirty-seven out of the 48 screened mutants showed the “striated” cell arrangement phenotype of the WT (**Figure 3A, B**). Interestingly this class contains several categories of mutants that show altered biofilm development in diverse models [63,67]. These include mutants with altered levels of second messengers; those with defects in primary metabolism; those with deletions of oxygen-sensing or NO-sensing proteins; and those with defects in the production of flagellar components, fimbriae, Pel polysaccharide, or selected virulence factors. This finding shows that effects on biofilm formation and development at the macroscale do not correlate with effects on cellular arrangement at the microscale and across depth.

The remaining 11 mutants showed cellular arrangement phenotypes that differed from the WT. The “bundled” phenotype was observed in all tested genetic knockouts defective in the production and function of the pilus (i.e., *pil* gene mutants and the mutant lacking RpoN, which is required for *pilA* expression) [68,69] (**Figure 3A, B** and **Figure S1**). In this phenotype, the pattern of cellular orientation is similar to that observed for the WT, but striations appear to be bundled because the fluorescently labeled populations are wider and show broader spacing between each other than WT striations. We hypothesize that pilus-mediated motility, known as twitching, serves to distribute cells in the x-y plane before vertical growth in zone 2 promotes the formation of segregated striations in the WT. According to this model, bundled striations form in twitching-defective mutants because daughter cells are less able to move away from their relatives before the elongation of zone 2.

Biofilms with the “disordered” phenotype display a “zone 1”-type distribution of cells across their full depth, i.e. cells are evenly spaced and randomly oriented (**Figure 3B, Figures S1** and **S3A**). This was observed in mutants with disruptions in the GacS/GacA two-component system or LasR, both of which constitute regulators that affect the expression of hundreds of genes. In addition, deletion of the O-antigen biosynthetic gene *wbpM* gave rise to this phenotype (**Figure 3B**). O-antigen is the outermost portion of lipopolysaccharide (LPS), a major constituent of the outer membrane of Gram-negative bacteria [70] (**Figure 5A**). The appearance of these biofilms suggests that in these mutants, related cells are fully dispersed from each other. Finally, Δ*ssg* and Δ*wapR*–mutants that are predicted to form LPS without the O-antigen attached (**Figure 3B** and **Figure S1**) [71–73]--gave rise to a “clustered” phenotype. These biofilms showed random cell orientation, but fluorescent cells also formed clusters with vertical elongation in zones 2 and 3, indicating that these mutants retained some ability to arrange related cells along the y-axis (**Figure 3B** and **Figure S1**). We note that our screen did not yield any mutants that exhibit vertical cellular orientation without clustering of related cells (or striation formation). This may indicate that, in zone 2, dispersion after cell division overrides any ability to establish vertical orientation.

### A retractable pilus is required for WT cellular arrangement across biofilm depth

To gain further insight into the structure of the bundled phenotype, which arises in mutants with pilus-related defects (**Figure 3A, B** and **4A**), we used high-resolution light sheet microscopy to take top-view images. For this method, the medium was inoculated using a cell suspension containing the P_PA1/04/03_-mScarlet strain at 1% and nonfluorescent cells at 99%. Macrocolonies were imaged after 19 hours of growth. The WT and Δ*pilA* biofilms showed patterns of fluorescence consistent with the distribution of fluorescent cells we observe in biofilm thin-sections taken between 12 and 24 hours (**Figures 1C** and **3D**): striations appears as puncta or as vertical columns that are tilted at an angle away from perfect perpendicularity with the biofilm-air interface. Fluorescent striations in the Δ*pilA* biofilms are thicker and occur at greater distances away from each other than those in WT biofilms. In Δ*pilA*, the striations also generally show a vertical orientation in the biofilm center, but diagonal orientations that radiate toward the colony edge in the biofilm outgrowth (**Figure 4A**). This arrangement is reminiscent of the “hedgehog”-like cellular ordering that has been reported for *Vibrio cholerae* biofilms grown from single cells under flow [12,14].

**Figure 4.**
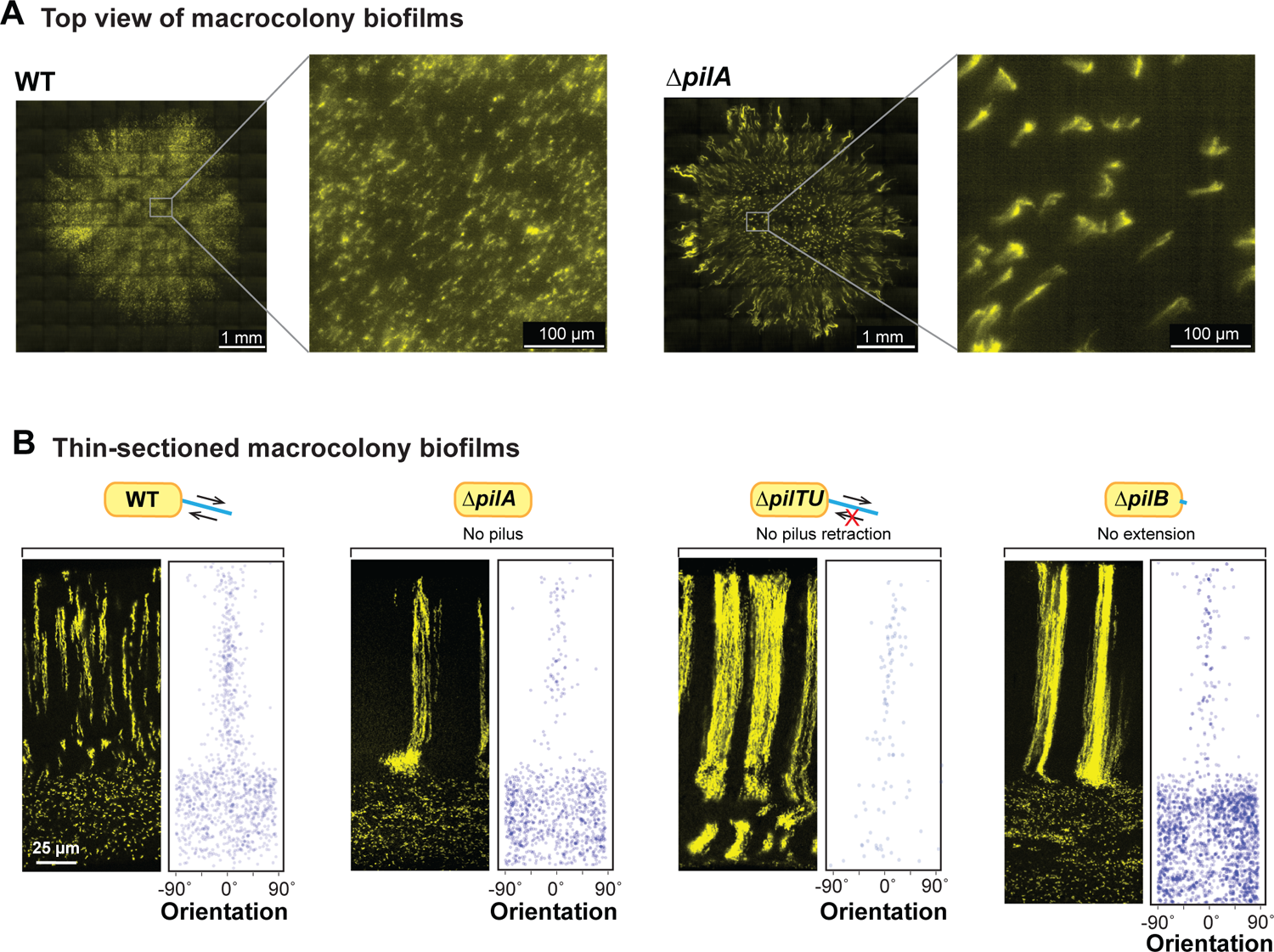
A retractable pilus is required for WT cellular arrangement. (**A**) High-resolution light sheet microscopy images of 19-hour old biofilms, with 1% of cells expressing fluorescent protein (colored yellow). (**B**) Fluorescence micrograph and quantification of orientation for WT and each of the indicated mutants grown in the mixing assay and thin-sectioned. The extent of pilus function present in each strain is indicated by a cartoon.

The phenotypes of the individual *pil* mutants allow us to make inferences about the contributions of the pilus and twitching motility–to cellular arrangement in WT biofilms. Twitching motility is conferred via repeated pilus extension, attachment, and retraction. The primary constituent of the pilus is PilA, also known as pilin. PilB is the extension motor–responsible for pilus assembly–while PilTU is the retraction motor [74,75]. PilO and PilY1 are also both required for pilus assembly; moreover, recent studies have revealed roles for both PilY1 and PilT in surface sensing [76–78]. All of the *pil* mutants we tested show bundled striations in zone 2 highlighting that a functional, i.e., retractable pilus is required for the formation of wild-type striations (**Figure 4B** and **S1**); however, Δ*pilTU* is unique in that it also shows bundled striations in zone 1 (**Figure 4B**).

### PilA is required for the zone-2 dispersion observed in O-antigen synthesis and attachment mutants

The results of our screen provide clues regarding the molecular determinants of cellular arrangement in *P. aeruginosa* macrocolonies. First, they suggest that a functional pilus is required for cellular distribution specifically and immediately before the formation of zones 2/3, i.e. that related cells twitch away from each other before they divide and form vertical lineages. However, the pilus is not required for cellular dispersion in zone 1, vertical striation formation, or vertical orientation (**Figure 3B** and **4A, B**). Second, they suggest that modifications to the structure of LPS can affect both cellular orientation and the clustering of related cells (or striation formation). These inferences raise the question of whether these cell surface components show interactions at the regulatory or phenotypic level. We tested this by evaluating PilA protein levels in O-antigen synthesis/attachment mutants and by examining cellular arrangement in combinatorial mutants.

To assess the effect of the LPS mutants on cellular PilA levels, we grew them as macrocolony biofilms for three days before sample preparation. PilA levels were comparable in all tested strains, WT, Δ*wbpM*, Δ*wapR*, and Δ*ssg*, indicating that the distinct cell-arrangement phenotypes are not due to differences in total PilA levels (**Figure 5B**).

**Figure 5.**
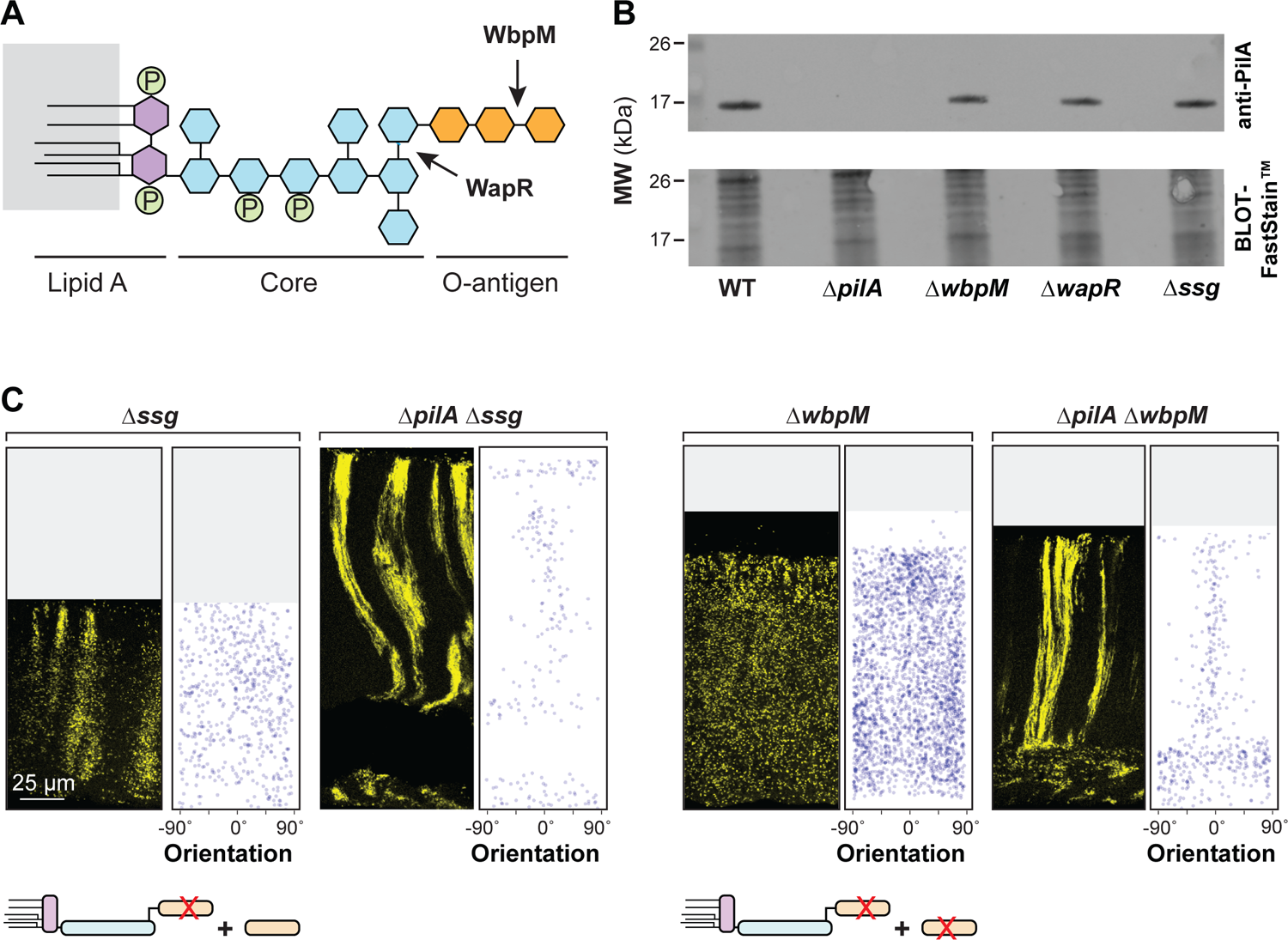
PilA is required for the disordered cellular arrangement phenotypes of O-antigen mutants. (**A**) Schematic of LPS and O-antigen indicating the bonds affected by WapR and WbpM activity. Hexagons represent monosaccharides and circles represent phosphate groups. The major components of LPS are color-coded. (**B**) Western blot showing PilA protein levels in macrocolony biofilms of WT and the indicated mutants. Equal amounts of total protein from sheared whole-cell lysates for each strain were resolved by SDS-PAGE using a 15% polyacrylamide gel. The PilA protein was detected using an anti-PilA antibody. (**C**) Fluorescence micrograph and quantification of orientation for each of the indicated mutants grown in the mixing assay and thin-sectioned. A cartoon representation of LPS and O-antigen (colors corresponding to panel (**A**)) indicates whether unattached and/or attached O-antigen are present in each strain. Images shown are representative of at least two independent experiments and mScarlet fluorescence is colored yellow.

To examine phenotypic interactions between mutations that affect O-antigen synthesis/attachment and that which affects pilin production, we created double mutants lacking combinations of the genes *wbpM*, *ssg*, and *pilA*, both with and without the P_A1/04/03_-*mScarlet* construct. Macrocolony biofilm mixing assays revealed that Δ*wbpM* is epistatic on Δ*ssg* (**Figure S1**). This indicates that the presence of O-antigen that is not attached to LPS in Δ*ssg* is sufficient to promote the clustering of related cells observed in this single-gene mutant.

Intriguingly, they also revealed that Δ*pilA* is epistatic on Δ*wbpM* and Δ*ssg*: deleting *pilA* in these backgrounds restored the formation of (bundled) striations (**Figure 5C**). These observations suggest that *pilA* promotes dispersion in both the striated WT and non-striated mutants, such as Δ*ssg* and Δ*wbpM*. In the absence of O-antigen-modified LPS, *pilA* also appears to promote random cellular orientation. This finding highlights the intricate interplay between pilus function, O-antigen synthesis/attachment, and cellular organization in the construction of *P. aeruginosa* biofilms. Interestingly, although it remains unconfirmed in *P. aeruginosa* PA14, pilus glycosylation with O-antigen has been described in other *P. aeruginosa* strains [79]. In this context it is also noteworthy that LPS-dependent changes in hydrophobicity have been shown to correlate with altered cellular packing in *P. aeruginosa* aggregates [80].

### Cell patterning influences the distribution of substrates across biofilm depth

The distribution of resources throughout a biofilm can be affected by a variety of factors, including its physical structure, properties of the matrix, and the metabolic activity of the cells within [21,81]. To examine whether cellular arrangement impacts how a substrate is distributed within *P. aeruginosa* macrocolonies, we engineered our strains of interest to contain a *rhaSR-PrhaBAD* inducible promoter system designed to express *mScarlet* in response to the small molecule L-rhamnose [82] (**Figure 6C**). Corresponding strains that constitutively produce eGFP were made so that related-cell distribution and rhamnose-induced expression could be examined in the same thin section. Macrocolony biofilms were grown from inocula that contained the constitutive eGFP-producing strain at 2.5% and the RhaSR-*PrhaBAD*-controlled mScarlet-producing strain at 97.5%. These were incubated for three days on our standard medium, then transferred to medium containing L-rhamnose for five hours before they were embedded in paraffin and prepared for thin sectioning (**Figure 6A**). Using this system, we found that in zones 1 and 2 mScarlet production was undetectable or only slightly above that observed in uninduced controls for all strains, indicating that pleiotropic effects inhibited activity of the *rhaSR-PrhaBAD* inducible promoter system in this part of the biofilm. However, all strains showed production of mScarlet in zone 3, i.e. at the biofilm-air interface. We will therefore focus our interpretations only on zone 3. WT, Δ*pilA,* and Δ*pilA*Δ*wbpM* biofilms showed similar, moderate levels of mScarlet production in this region. In the two disordered mutants Δ*wbpM* and Δ*gacA* (**Figure 6B, D** and **Figure S2**), mScarlet production in this region was greatly enhanced. We note that deletion of *wbpM* is not expected to independently affect *PrhaBAD* activity [83]. These findings suggest that lengthwise cellular packing and/or vertical cellular orientation affect the transport of nutrients across biofilm depth.

**Figure 6.**
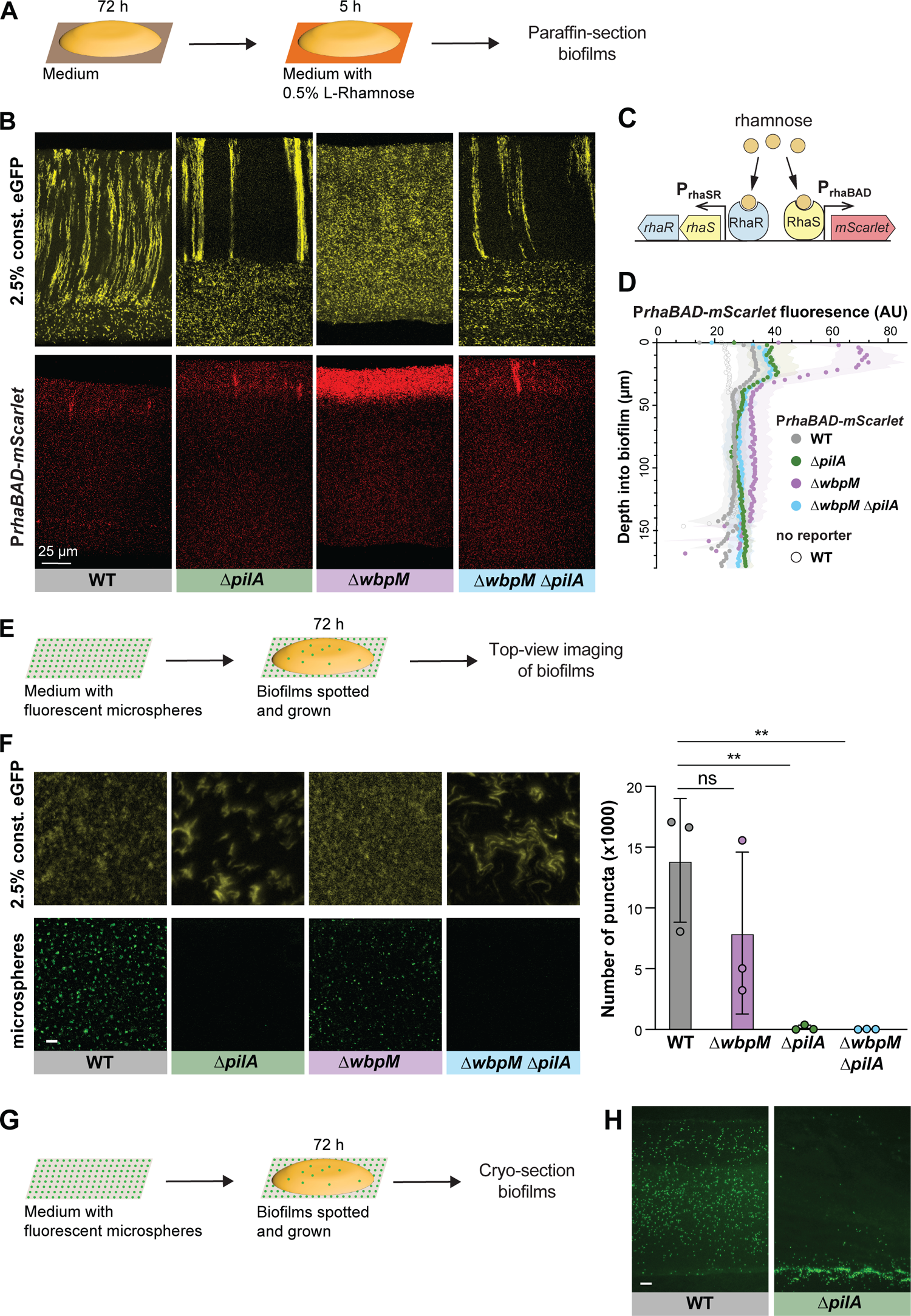
Cellular arrangement affects the uptake of substrates into colony biofilms. (**A**) Schematic illustration of the experimental setup for growing *P. aeruginosa* biofilms on agar plates and their subsequent transfer to medium containing L-Rhamnose. (**B**) Fluorescence micrographs of thin sections from WT and indicated mutant biofilms. Biofilm inocula contained 2.5% cells that constitutively express eGFP and 97.5% RhaSR-*PrhaBAD*-controlled mScarlet-producing strain. Top panels show eGFP fluorescence (colored yellow) and bottom panels show the mScarlet fluorescence for each thin section. (**C**) Schematic of RhaSR-P*rhaBAD* expression system driving mScarlet production. (**D**) Quantification of mScarlet fluorescence shown in (B). Shading represents standard deviation of biological triplicates. (**E**) Schematic of the experimental setup for growing *P. aeruginosa* macrocolony biofilms on agar plates with fluorescent microspheres (200 nm). (**F**) **Left**: Top-view images taken from center of macrocolony biofilms that were grown on medium with microspheres (colored green). Biofilms contained 2.5% cells constitutively expressing eGFP (colored yellow). **Right**: Quantification of microspheres visible in top-view images. Each data point is a biological replicate, bar height indicates the mean of these replicates. p values are based on two-sided unpaired t-tests (n.s., not significant; ** p ≤ 0.01) Scale bar is 25 µm. (**G**) Schematic illustration of the experimental setup for growing biofilms on agar plates with fluorescent microspheres for cryosectioning. (**H**) Fluorescence micrographs of cryosections of biofilms grown on microspheres (colored green). Scale bar is 25 µm.

To further examine the effect of cellular arrangement on transport within biofilms, we tested the uptake and distribution of fluorescent microspheres (diameter: 200 nm). Microspheres were spread on the surface of an agar plate before inoculation for macrocolony biofilm growth. After a three-day incubation, the biofilms were imaged from the top or cryosectioned and imaged immediately (**Figure 6E, G**). The microspheres were visible at the air-exposed surface of WT and Δ*wbpM* biofilms. However, in the absence of type IV pili (ΔpilA and ΔpilA ΔwbpM) the microspheres were barely able to pass through to the biofilm surface (**Figure 6F**), indicating pilus-dependent changes in cellular arrangement. To probe the distribution of microspheres within biofilms, we cryosectioned the WT (spheres visible at the surface) and the Δ*pilA* mutant (spheres not visible at the surface). Indeed, microspheres were evenly distributed within WT biofilms, while for Δ*pilA* they were barely able to enter the biofilm at the agar interface (**Figure 6H**). In summary, the distribution of substrates of different sizes (rhamnose and microspheres)--into and within biofilms–was affected by changes in cell arrangement.

### Altered cellular arrangement phenotypes correlate with effects on the distributions of metabolically active and surviving cells

The correlations we observed between cellular-arrangement phenotypes (**Figure 3B**), metabolic activity profiles (**Figure 2B**), and substrate distribution (**Figure 6B, F, H and Figure S2**) in biofilms suggested that mutations that affect microorganization across depth could affect physiological patterning. We therefore tested whether mutant biofilms with altered cellular arrangement show altered distributions of metabolic activity (**Figure 7A**). SRS imaging of mutant biofilm thin-sections indeed revealed that the band of metabolic activity is shifted closer to the top of the biofilm in Δ*wbpM* (peak “2”) when compared to WT (peak “3”) and is unchanged in the Δ*pilA* mutant. The metabolic activity profile of Δ*wbpM* also shows a “shoulder” of increased activity (peak “1”) that is not present in the WT and Δ*pilA*. A double deletion of *wbpM* and *pilA* lacked this shoulder (**Figure 7B**).

**Figure 7.**
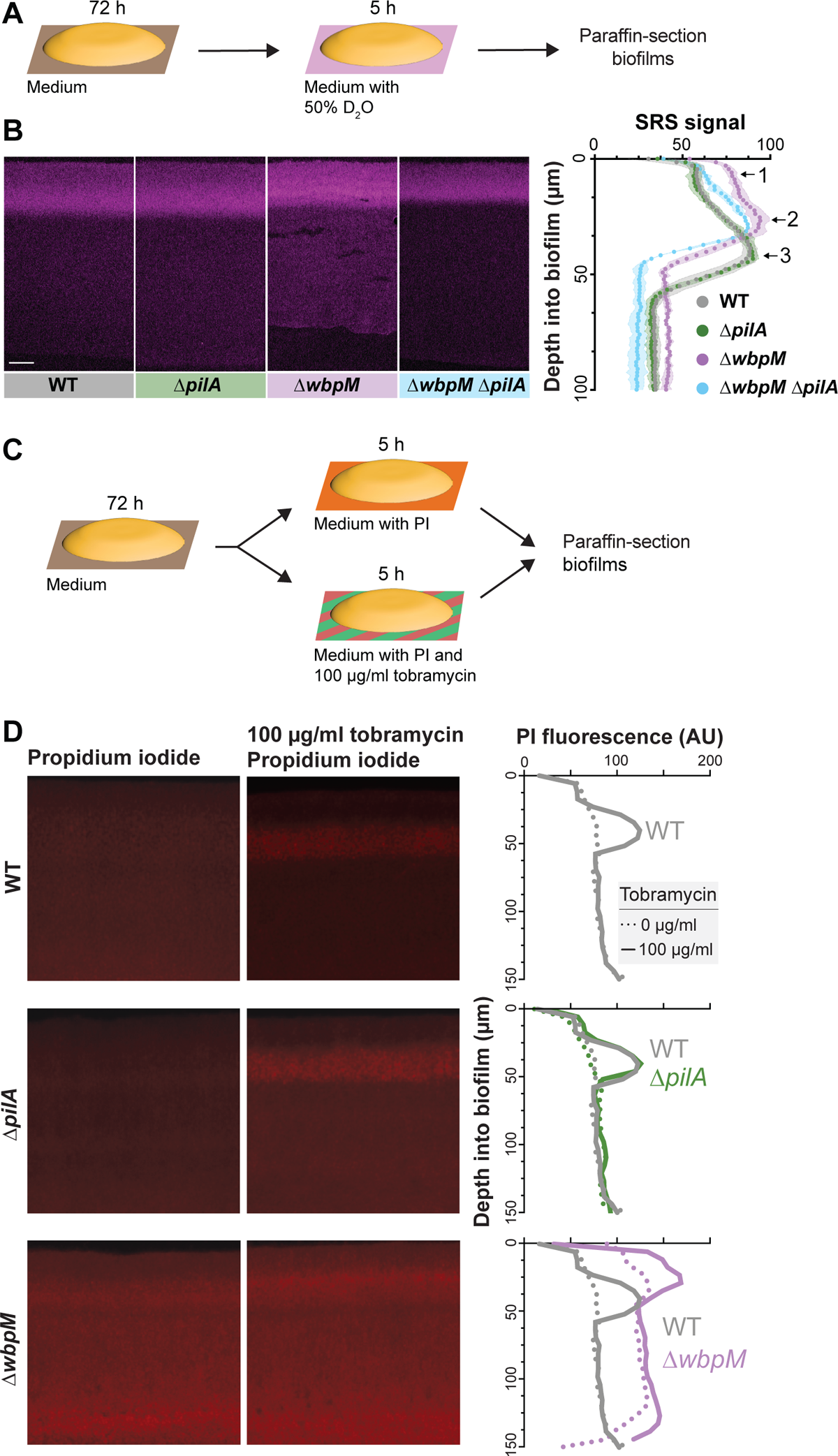
Mutations affecting cellular arrangement alter metabolic activity and antibiotic tolerance profiles within biofilms. (**A**) Schematic illustration of the experimental setup for growing biofilms on agar plates and their subsequent transfer to a medium containing D_2_O for analysis of metabolic activity using SRS microscopy. (**B**) **Left**: SRS micrographs of WT and mutant biofilm thin-sections. Scale bar, 25 µm. **Right**: Graph showing the quantitative shift in metabolic activity distribution in different mutants. (**C**) Schematic illustration of the experimental setup for growing biofilms on agar plates and their subsequent transfer to a medium containing propidium iodide or propidium iodide + tobramycin (**D**) **Left**: Fluorescence micrographs of three-day-old biofilms exposed to propidium iodide (PI), a DNA stain. **Center**: Fluorescence micrographs of biofilms treated with tobramycin and PI. **Right**: Graphs showing quantification of PI staining. Images are representative of at least two independent experiments.

To further interrogate physiological status, we assessed cell death across depth in biofilms by exposing them to propidium iodide (PI), a DNA stain that is excluded from bacteria with intact membranes. For this assay, three-day-old biofilms were transferred to plates containing medium supplemented with 50 µM propidium iodide, and incubated for five hours before sample preparation and thin-sectioning (**Figure 7C**). Microscopic imaging revealed increased PI staining in thin sections of Δ*wbpM* biofilms, particularly in the upper region of the biofilm, when compared to WT. Thin sections of WT and Δ*pilA* biofilms showed similar levels and distributions of PI staining (**Figure 7D**).

Together, the results of our metabolic activity and cellular survival profiling experiments are consistent with the notion that more substrates from the growth medium reach the biofilm region at the air interface in the Δ*wbpM* mutant. The altered cellular arrangement in the Δ*wbpM* mutant biofilm appears to disrupt the normal stratification of metabolic activity, pushing it closer to the biofilm surface (**Figure 7B**). The increased PI staining in this region could be a consequence of this accelerated metabolic activity, leading to quicker exhaustion of local resources and, thus, a higher incidence of non-viable cells.

### Altered cellular-arrangement patterns correlate with decreased antibiotic tolerance

The physiological status of a bacterium affects its susceptibility to antibiotics, and particularly those that target biosynthetic reactions and/or that require functional membrane transport processes for uptake [84–87]. Because we had observed that mutants with altered cellular arrangements showed distinct patterns of substrate distribution and metabolic activity within biofilms (**Figure 6B, F, H, Figure 7B, and Figure S2**), we hypothesized that they would be differentially susceptible to antibiotic treatment. To test this, we transferred three-day-old *P. aeruginosa* macrocolonies to a medium containing propidium iodide and tobramycin, an aminoglycoside antibiotic that requires proton motive force for its bactericidal action, and incubated for five hours before biofilms were prepared for thin-sectioning [88]. We infer that an increase in propidium iodide staining indicates a decrease in antibiotic tolerance [89,90].

Comparison to thin-sections from biofilms that had not been treated with tobramycin (**Figure 7D**) revealed that, in WT and Δ*pilA*, the antibiotic killed cells in the region spanning from 20-60 µm from the air-biofilm interface, which corresponded to the regions of high metabolic activity (**Figure 7B**). In contrast, Δ*wbpM* and Δ*gacA* biofilm thin-sections showed tobramycin-dependent cell death in a region that was closer to the air-biofilm interface (0-40 µm depth) (**Figure 7D, Figure S3B, S3C**), corresponding to the shifted region of high metabolic activity that we had observed for Δ*wbpM* (**Figure 7B**). In agreement with the dominant effect of the *pilA* deletion on cell arrangement (**Figure 5C** and **Figure S3A**) and metabolic activity (**Figure 7B**), the peak of propidium iodide staining shifted back to a depth of 20-60 µm in the Δ*gacA*Δ*pilA* double mutant (**Figure S3C**). While cells are randomly oriented in Δ*wbpM* and Δ*gacA* biofilms, the deletion of *pilA* in these backgrounds results in vertical cellular arrangement as observed in WT and Δ*pilA* (**Figure 5C** and **Figure S3A**). These patterns further highlight the correlation between cellular arrangement and antibiotic tolerance.

Because our data suggest that cellular arrangement affects substrate transport across biofilm depth, two phenomena may be contributing to the differential impact of tobramycin treatment on ordered (WT) and disordered-mutant biofilms. First, tobramycin, like rhamnose, may move more readily through disordered biofilms than through ordered ones, leading to a shift in the zone of tobramycin-dependent cell death toward the biofilm-air interface. Alternatively or in addition, the greater availability of rhamnose near the biofilm-air interface in disordered mutants like Δ*wbpM*– which appears to enhance metabolic activity in this zone–may also contribute to tobramycin susceptibility. Regardless of whether one or both of these phenomena are responsible for changes in the disordered-mutant profiles, our data indicate a relationship between cellular organization and the physiological properties of the biofilm that impacts antibiotic tolerance.

### Concluding remarks

This study has uncovered genetic and structural determinants that orchestrate cell arrangement within *P. aeruginosa* biofilms. A screen of 48 mutants representing genes implicated in biofilm development and physiology identified 11 that impact cellular organization across depth, underscoring that this aspect of biofilm architecture is uniquely controlled. Our work also revealed correlations between cellular arrangement phenotypes and impacts on substrate distribution, metabolic activity, and antibiotic tolerance highlighting the critical role of microorganization in determining metabolic status. An understanding of the relationships between microanatomy and physiology, not only deepens our knowledge of bacterial multicellularity and provides avenues for the development of new therapeutic strategies. It also prompts further exploration into the potential parallels and divergences between biofilm mechanisms and multicellular organization in other organisms, paving the way for interdisciplinary insights.

## MATERIALS AND METHODS

### Bacterial strains and culture conditions

Bacterial strains used in this study are listed in Table 1. Liquid cultures were routinely grown in Lysogeny Broth (LB) [91] at 37 °C with shaking at 250 rpm.

### Construction of mutant *P. aeruginosa* strains

To create markerless deletion mutants in *P. aeruginosa* PA14, 1 kb of flanking sequence from each side of the target gene were amplified using the primers listed in Table 3 and inserted into pMQ30 through gap repair cloning in Saccharomyces cerevisiae InvSc1 [92]. Each plasmid listed in Table 2 was transformed into Escherichia coli strain UQ950, verified by sanger sequencing, and moved into PA14 using biparental conjugation. PA14 single recombinants were selected on LB agar plates containing 100 µg/ml gentamicin. Double recombinants (markerless deletions) were selected on LB without NaCl and modified to contain 10% sucrose. Genotypes of deletion mutants were confirmed by PCR. Combinatorial mutants were constructed by using single mutants as hosts for biparental conjugation.

### Scanning electron microscopy of macrocolonies

Overnight precultures were diluted 1:100 in LB and grown to mid-exponential phase (OD at 500 nm ∼ 0.5). OD values at 500 nm were read in a Spectronic 20D+ spectrophotometer (Thermo Fisher Scientific [Waltham, MA]) and cultures were adjusted to the same OD. Five microliters of mixture were spotted onto a bilayer plate. After three days of incubation, macrocolonies were carefully overlaid with 1% agar to preserve their structure. Blocks were then cut out, and the bottom layer of the agar was carefully removed, leaving an equal volume on top and below the macrocolony. The blocks were then placed on filter paper to facilitate moving them when necessary with the least amount of disturbance and placed into an empty glass petri dish. The blocks were then incubated in McDowell Trump Fixative for 16 hours at room temperature to allow fixation to occur. They were washed twice with 0.1 M PBS before being subjected to 1% (wt/vol) Osmium Tetroxide in 0.1 M PBS to further fix the samples. The samples were then washed twice with distilled water before being subjected to dehydration in an increasing graded ethahol series (35%, 50%, 75%, 2X 90%, 3X 100%). The blocks were washed twice in HMDS before being mounted onto aluminum stubs using double-sided round carbon stickers and placed into a dessicator. After air-drying thoroughly, the blocks were cut in half in order to visualize the internal structure and orientation of cells within the biofilm. The samples were then coated in gold using a Leica EM ACE600 Coater. The samples were visualized using an Helios Nanolab DualBeam 660 (FEI) operating at an accelerating voltage of 5 kV under a high vacuum.

### Colony biofilm mixing assay

Overnight precultures were diluted 1:100 in LB and grown to mid-exponential phase (OD at 500 nm ∼ 0.5). OD values at 500 nm values were read in a Spectronic 20D+ spectrophotometer (Thermo Fisher Scientific [Waltham, MA]) and cultures were adjusted to the same OD. Adjusted cultures were then mixed in a 2.5:97.5 ratio of fluorescent:non-fluorescent cells. Five microliters of mixture were spotted onto a bilayer plate of 45 mL (bottom layer) and 15mL (top layer) of 1% tryptone, 1% agar [Teknova (Hollister, CA) A7777] in a 10 cm x 10 cm x 1.5 cm square Petri dish (LDP [Wayne, NJ] D210-16). Plates were incubated for three days at 25°C in the dark and imaged using a VHX-1000 digital microscope (Keyence, Japan).

### Paraffin embedding, thin sectioning and confocal imaging

After three days of growth as described above, biofilms were overlaid with 1% agar and sandwiched biofilms were lifted from the bottom layer and fixed overnight in 4% paraformaldehyde in PBS at 25 °C for 24 h in the dark. Fixed biofilms were washed twice in PBS and dehydrated through a series of 60-min ethanol washes (25%, 50%, 70%, 95%, 3 × 100% ethanol) and cleared via three 60-min incubations in Histoclear-II (National Diagnostics); these steps were performed using an STP120 Tissue Processor (Thermo Fisher Scientific).

Biofilms were then infiltrated with wax via two separate 2-h washes of 100% paraffin wax (Paraplast Xtra) at 55 °C, and allowed to polymerize overnight at 4 °C. Trimmed blocks were sectioned in 10-µm-thick sections perpendicular to the plane of the biofilm, floated onto water at 42 °C, and collected onto slides. Slides were air-dried overnight, heat-fixed on a hotplate for 30 minutes at 45 °C, and rehydrated in the reverse order of processing. Rehydrated colonies were immediately mounted in (Thermo Fisher Scientific ProLong™ Diamond Antifade Mountant P36965) and overlaid with a coverslip. Fluorescent images were captured using an LSM800 confocal microscope (Zeiss, Germany). Samples were illuminated with 488 and 561 nm lasers, respectively. eGFP acquisition parameters were: 400-548nm (emission wavelength range). mScarlet acquisition parameters were: 593-700nm (emission wavelength range). Each strain was prepared in this manner in at least biological triplicates.

### Image analysis and quantification of biofilm microorganization

Image analysis was performed in FIJI [93] and Python (Python Software Foundation; Python Language Reference, version 3.8. Available at www.python.org). Each image was viewed in FIJI for quality control and then binarized using a custom Python script. For each sample, a fixed-threshold binarization was applied to the fluorescence image using a multi-level version of Otsu’s method, which separates pixels of an input image into several classes obtained according to intensity [94]. For most images, the number of classes *n* was set to three, and the image was binarized by setting pixels in the brightest class equal to one and all others to zero. Qualitative inspection of the thresholded images were performed for quality assurance; samples which yielded poor separation into three classes (due to, e.g., lower resolution or fluorescence intensity) were binarized using the classic Otsu’s method [95]. Images were then cleaned by using masks to remove artifacts. All successive analysis and quantification was performed using the cleaned binarized images.

Boundaries of individual cells were segmented from binarized images using the Python SciPy package [96]. The orientation of each cell was determined as the angle between the horizontal and the major axis of the best-fit ellipse to each cell, ranging from −90 to +90 degrees. Heights along the biofilm’s direction of growth were divided into bins of 25 pixels in width. An order parameter to quantify cell alignment was defined as the standard deviation of the orientation of all cells within a bin; a high value indicates that cells in a bin are randomly oriented, while a low value indicates that cells are aligned in the same direction. All code is available at https://github.com/jnirody/biofilms.

### Pellicle biofilm mixing assay

Overnight precultures of fluorescent (mScarlet-expressing) and non-fluorescent strains were diluted 1:100 in LB and grown to mid-exponential phase (OD at 500 nm ∼ 0.5). OD values at 500 nm values were read in a Spectronic 20D+ spectrophotometer (Thermo Fisher Scientific [Waltham, MA]) and cultures were adjusted to the same OD. Adjusted cultures were then mixed in a 2.5:97.5 ratio of fluorescent:non-fluorescent cells and nine ml of this mixture was added to 13×100mm culture tubes (VWR, Cat no. 10545-936) and grown in 25°C incubator for three days. Pellicles were removed from glass tube using a sterile needle and transferred to plate and overlayed with 1% agar. Samples were subsequently processed using standard paraffin embedding and thin sectioning protocol.

### Stimulated Raman Scattering Microscopy

Five microliters of a bacterial subculture were spotted onto 45mL/15mL two-layer 1% tryptone 1% agar plates and were grown for 72 hours, then the top layer and biofilm was transferred to 2 mL 1% tryptone 1% agar 50% D_2_O and incubated at 25°C for 5 hours or 24 hours. The biofilms were then thin sectioned as described above. For SRS microscopy, an integrated laser source (picoEMERALD, Applied Physics & Electronics, Inc.) was used to produce both a Stokes beam (1064nm, 6ps, intensity modulated at 8MHz) and a tunable pump beam (720–990nm, 5–6ps) at an 80MHz repetition rate. The spectral resolution of SRS is FWHM=6–7cm^−1^. These two spatially and temporally overlapped beams with optimized near-IR throughput were coupled into an inverted multiphoton laser-scanning microscope (FV1200MPE, Olympus). Both beams were focused on the biofilm samples through a 25X water objective (XLPlan N, 1.05 N.A. MP, Olympus) and collected with a high N.A. oil condenser lens (1.4 N.A., Olympus) after the sample. By removing the Stokes beam with a high O.D. bandpass filter (890/220 CARS, Chroma Technology), the pump beam is detected with a large area Si photodiode (FDS1010, Thorlabs) reverse-biased by 64 DC voltage. The output current of the photodiode was electronically filtered (KR 2724, KR electronics), terminated with 50Ω, and demodulated with a lock-in amplifier (HF2LI, Zurich Instruments) to achieve near shot-noise-limited sensitivity. The stimulated Raman loss signal at each pixel was sent to the analog interface box (FV10-ANALOG, Olympus) of the microscope to generate the image. All images were acquired with 80 µs time constant at the lock-in amplifier and 100µs pixel dwell time (∼27s per frame of 512×512 pixels). Measured after the objectives, 40mW pump power and 120mW Stokes beam were used to image the carbon-deuterium 2183 cm^-1^ and off-resonance 2004 cm^-1^ channels.

### Macrocolony clearing and light sheet microscopy

Overnight precultures were diluted 1:100 in LB and grown to mid-exponential phase (OD at 500 nm ∼ 0.5). OD values at 500 nm values were read in a Spectronic 20D+ spectrophotometer (Thermo Fisher Scientific [Waltham, MA]) and cultures were adjusted to the same OD. Adjusted cultures were then mixed in a 1:100 ratio of fluorescent:non-fluorescent cells. 0.5 microliters of mixture were spotted onto a bilayer plate of 50 mL (bottom layer) and 10 mL (top layer) of 1% tryptone, 1% agar [Teknova (Hollister, CA) A7777] in a 10 cm x 10 cm x 1.5 cm square Petri dish (LDP [Wayne, NJ] D210-16). Plates were incubated for one day at 25°C in the dark.

Biofilms were overlaid with 1% agar, and sandwiched biofilms were lifted from the bottom layer, and fixed overnight in 4% paraformaldehyde in PBS at 25 °C for 12 hours in the dark. Fixed biofilms were washed twice in PBS then incubated for three hours in 70% (V/V) glycerol solution deionized water solution for optical clearing. The clarified biofilms were mounted on a standard microscopy slide, and were imaged with ClearScope [97] (MBF Biosciences) light sheet microscope, using Nikon 20x/1.0NA detection objective. The resulting image tiles were stitched by using a Terastitcher based custom pipeline [98,99], and analyzed using ImageJ after 2-fold down-sampling in x-y axes.

### SDS-page and western

Sheared pilin and cell lysate proteins were extracted as previously described [100] with some modifications. Overnight precultures were diluted 1:100 in LB and grown to mid-exponential phase (OD at 500 nm ∼ 0.5). OD values at 500 nm values were read in a Spectronic 20D+ spectrophotometer (Thermo Fisher Scientific [Waltham, MA]) and cultures were adjusted to the same OD. Five microliters of mixture were spotted onto a plate of 60 mL of 1% tryptone, 1% agar [Teknova (Hollister, CA) A7777] in a 10 cm x 10 cm x 1.5 cm square Petri dish (LDP [Wayne, NJ] D210-16). Plates were incubated for three days at 25°C in the dark.

Biofilm samples (25 biofilms per strain) were gently scraped from agar, resuspended in 1 mL sterile PBS (phosphate buffered saline, pH 7.4), and centrifuged at 4000 x g for five minutes at room temperature. Pellets were resuspended in 4.5 mL of sterile PBS, vortexed for 30 s to shear cell surface proteins, and centrifuged at 4000 x g for 15 min at room temperature.

Supernatants containing the sheared surface proteins were subjected to multiple rounds of centrifugation at 11,600 x g for 15 min to remove residual cells from the samples. Next, 5 M NaCl and 30% (w/v) PEG (polyethylene glycol, ∼8000 MW, Sigma) at a volume ratio of 0.1 each were added to the total supernatant, and incubated on ice for 90 min before pilin was collected by centrifugation at 11,600 x g for 30 min at 4 °C. The pelleted pilin was resuspended in 1X LDS sample buffer (Invitrogen) with 5% β-mercaptoethanol (Sigma) and boiled for 5 min before separation on a 4-12% NuPAGE Bis-Tris gel (Invitrogen) for immunoblotting.

Cellular proteins were extracted from the cell pellets after surface pilin was sheared and removed (described above). Briefly, cell pellets were resuspended in 0.2 M Tris-Cl (pH 8.0), normalized to OD_600_ ∼ 4.0, centrifuged, and pellets were resuspended in 1X LDS sample buffer with 5% β-mercaptoethanol. Samples were boiled for five min, centrifuged, and aliquots from the supernatants were resolved on a 4-12% NuPAGE Bis-Tris gel for visualization and immunoblotting.

For PilA immunoblotting, both sheared and whole-cell pilin preparations were resolved using 4-12% NuPAGE Bis-Tris gels at 125V for 2.5 h and transferred onto 0.22 µm nitrocellulose membranes. Protein bands were normalized to total protein visualized by BLOT-FastStain (GBiosciences). Membranes were blocked using TBS-T (Tris-buffered saline with Tween-20) containing 5% milk before probing with an anti-PilA rabbit polyclonal antibody (gift of the O’Toole lab, Dartmouth) at 1:1000, followed by an IR conjugated goat anti-rabbit antibody at 1:10,000 (LI-COR Biosciences). Bands were visualized using the LI-COR Odyssey Clx imager and Image Studio.

### Rhamnose assay

Overnight precultures were diluted 1:100 in LB and grown to mid-exponential phase (OD at 500 nm ∼ 0.5). OD values at 500 nm values were read in a Spectronic 20D+ spectrophotometer (Thermo Fisher Scientific [Waltham, MA]) and cultures were adjusted to the same OD. Adjusted cultures were then mixed in a 2.5:97.5 ratio of constitutively fluorescent eGFP cells: rhamnose inducible mScarlet fluorescent cells. Five microliters of mixture were spotted onto bilayer plater. Plates were incubated for three days at 25 °C in the dark, then transferred to medium containing 0.5% L-rhamnose for five hours. Samples were subsequently processed using standard paraffin embedding and thin sectioning protocol.

### Microsphere assay

Overnight precultures were diluted 1:100 in LB and grown to mid-exponential phase (OD at 500 nm ∼ 0.5). OD values at 500 nm values were read in a Spectronic 20D+ spectrophotometer (Thermo Fisher Scientific [Waltham, MA]) and cultures were adjusted to the same OD. Adjusted cultures were then mixed in a 2.5:97.5 ratio of fluorescent:non-fluorescent cells. Bilayer plates of 45 mL (bottom layer) and 15 mL (top layer) of 1% tryptone, 1% agar [Teknova (Hollister, CA) A7777] in a 10 cm x 10 cm x 1.5 cm square Petri dish (LDP [Wayne, NJ] D210-16) were made and solidified overnight. 300 µl (10^10 microspheres/mL) of Fluoresbrite® Multifluorescent Microspheres 0.2 µm (PolySciences 24050-5) were spread onto a bilayer plate using sterile glass beads and dried for 30 minutes. Five microliters of bacterial culture mixture were spotted onto plate and incubated for three days at 25 °C in the dark and macrocolony biofilms were subsequently imaged live or prepared for cryosectioning. Samples were imaged on a ZEISS Axio Zoom.V16 microscope, equipped with a mercury lamp. eGFP was detected using a 450/490nm excitation filter, 500/550nm emission filter, and 495nm dichroic mirror. Fluoresbrite® Multifluorescent Microspheres were detected using a 538/562nm excitation filter, 570/640nm emission filter, and 570nm dichroic mirror.

### Macrocolony biofilm cryosectioning

Macrocolony biofilms were overlaid with 1% agar, and sandwiched biofilms were cut and lifted from the bottom layer, and placed in disposable cryomold (Tissue-Tek). They were then covered with an embedding agent (Tissue-Tek OCT), and flash frozen in liquid nitrogen. Cryoembedded biofilms were subsequently sectioned with a Leica CM1950 cryostat set at −18 °C. Ten-micrometer-thick sections perpendicular to the plane of the macrocolony were then transferred onto slides and immediately imaged using a ZEISS Axio Zoom.V16.

### Propidium iodide and antibiotic assay

Overnight precultures were diluted 1:100 in LB and grown to mid-exponential phase (OD at 500 nm ∼ 0.5). OD values at 500 nm were read in a Spectronic 20D+ spectrophotometer (Thermo Fisher Scientific [Waltham, MA]) and cultures were adjusted to the same OD. Adjusted cultures were then mixed in a 2.5:97.5 ratio of eGFP fluorescent:non-fluorescent cells. Five microliters of mixture were spotted onto bilayer plate. Plates were incubated for three days at 25 °C in the dark, then transferred to medium containing 50 µM PI with and without 100 µg/mL tobramycin for five hours. Samples were subsequently processed using standard paraffin embedding and thin sectioning protocol. Samples were imaged on a ZEISS Axio Zoom.V16 microscope, equipped with a mercury lamp. Propidium iodide staining was detected using a 538/562nm excitation filter, 570/640nm emission filter, and 570nm dichroic mirror.

## ACKNOWLEDGEMENTS

The authors thank Endah Rosa, Paula Fernandez Musso and Yu-Cheng Lin for assistance in generating mutants described in this manuscript. Thank you to the O’Toole Lab for gifting the anti-PilA rabbit polyclonal antibody. This work was supported by an NIH Diversity supplement to J.K. and by the parent NIH/NIAID grant, R01AI103369 to L.E.P.D. It was also supported by National Institute of Health (R01 EB029523) and Chan Zuckerberg Initiative (Dynamic Imaging 2023-321166) to W.M., a Rockefeller University Physics and Biology Fellowship and an All Souls College Fellowship in Life Sciences to J.A.N., NSF MCB 2216676 funding to A.J., and NIH DP2MH119423 and 1R44MH116827 funding to R.T.

