## Supplemental Files S1-S3 and Tables S1-S3 for "Cell arrangement impacts metabolic activity and antibiotic tolerance in *Pseudomonas aeruginosa* biofilms"

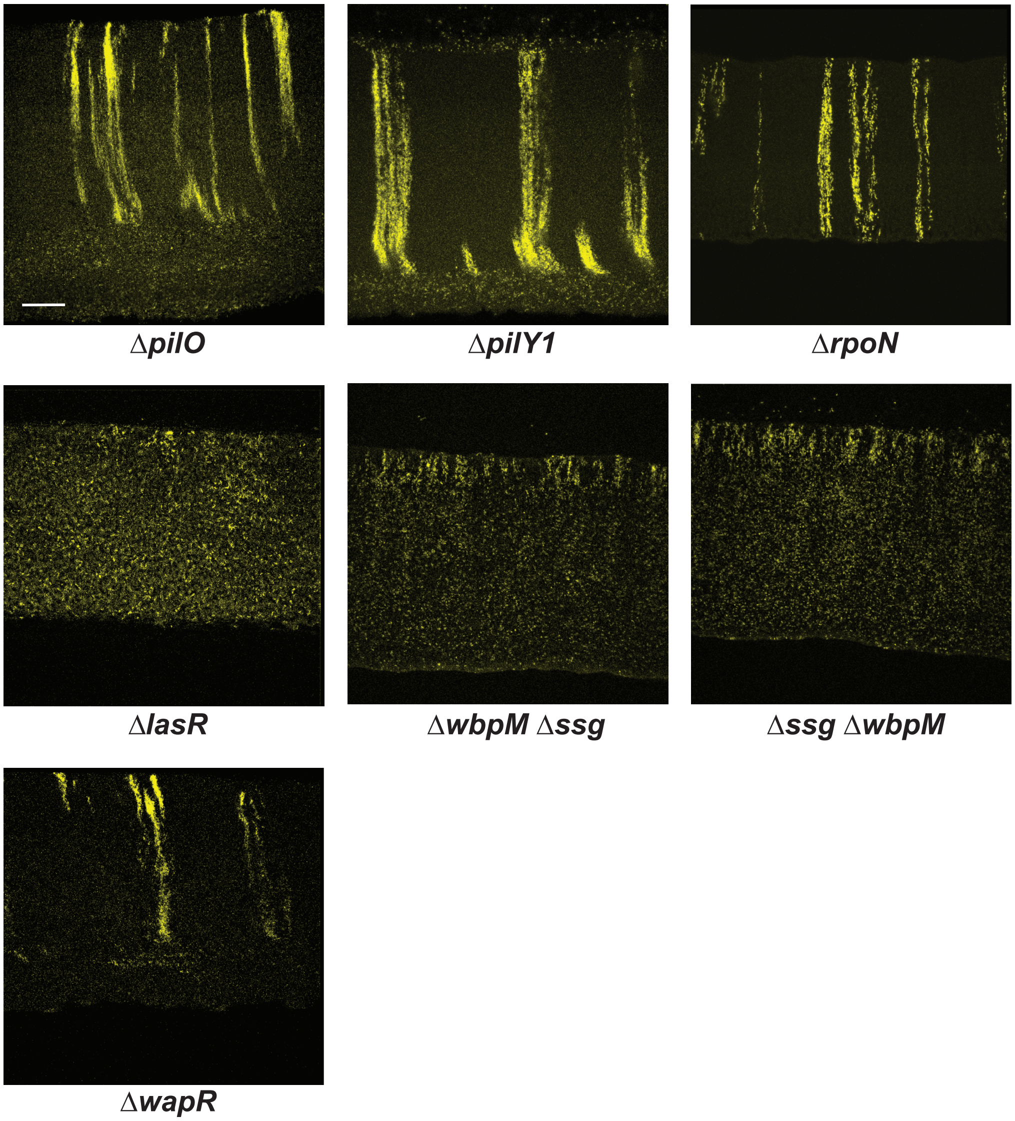


**Figure S1.Mutations affecting global regulators, pilus synthesis, and O-antigen synthesis/attachment alter cell patterning in colony biofilms.** Fluorescence micrographs of thin sections from indicated mutant biofilms grown on 1% tryptone and 1% agar for 3 days. The biofilm inocula contained 2.5% of cells that constitutively express mScarlet; 97.5% did not express a fluorophore. Scale bar, 25 μm. Images are representative of at least two independent experiments and mScarlet fluorescence is colored yellow.

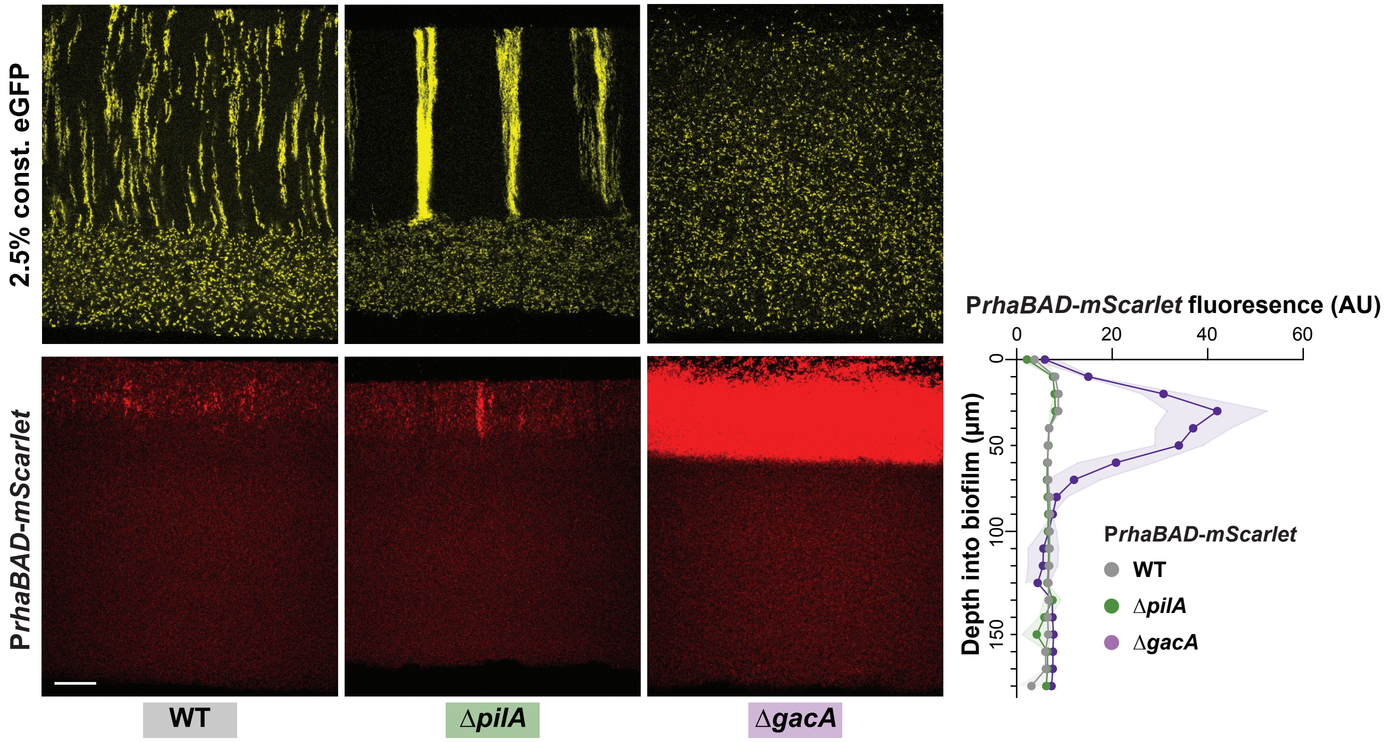


**Figure S2. Cellular arrangement affects the uptake and distribution of rhamnose into colony biofilms.** Fluorescence micrographs of thin sections from WT and indicated mutant biofilms. Biofilm inocula contained 2.5% cells that constitutively express eGFP and 97.5% RhaSR-*PrhaBAD*-controlled mScarlet-producing strain. Top panels show eGFP fluorescence (colored yellow) and bottom panels show the mScarlet fluorescence for each thin section. Scale bar, 25 μm. Quantification of mScarlet fluorescence is shown on the right.

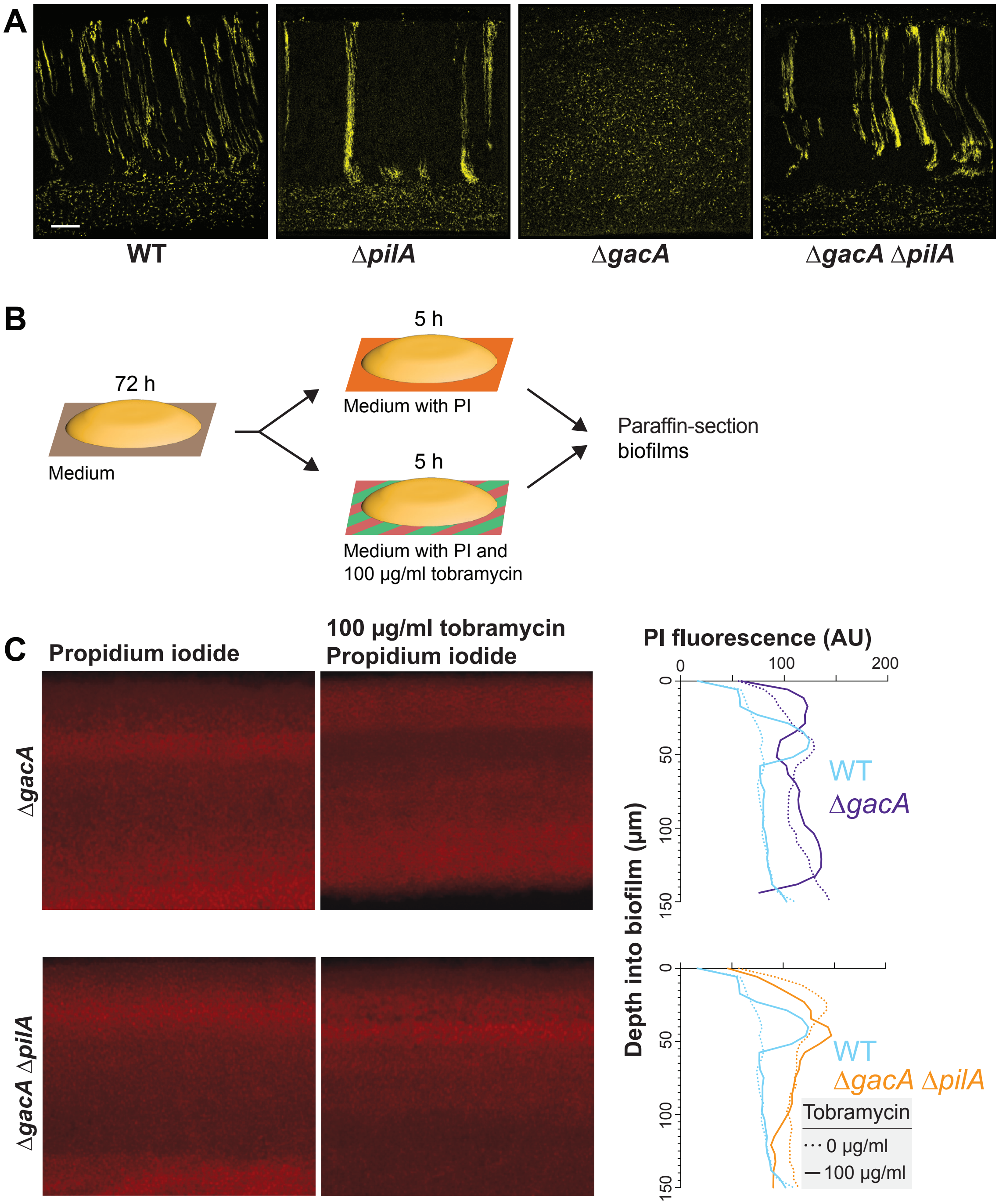


**Figure S3. Mutations affecting cellular arrangement alter biofilm antibiotic tolerance profiles.** (**A**) Fluorescence micrographs of thin sections from indicated mutant biofilms grown on 1% tryptone and 1% agar for three days. The biofilm inocula contained 2.5% of cells that constitutively express mScarlet (shown as yellow); 97.5% did not express a fluorophore. Scale bar, 25 µm. (**B**) Schematic illustration of the experimental setup for growing *P. aeruginosa* biofilms on agar plates and their subsequent transfer to a medium containing propidium iodide or propidium iodide + tobramycin (**C**) **Left**: Fluorescence micrographs of 3-day-old biofilms exposed to propidium iodide (PI), a DNA stain. **Center**: Fluorescence micrographs of biofilms treated with tobramycin and PI. **Right**: Graphs showing quantification of PI staining. Images shown in this figure are representative of at least two independent experiments.

**Table S1. Bacterial strains used in this study.**

| Number | Strain | Description | Source |
|---|---|---|---|
| *Pseudomonas aeruginosa* strains | | | |
| LD0 | UCBPP-PA14 (WT) | Clinical isolate UCBPP-PA14. | [1] |
| LD4058 | PA14 Δ*vfr* | PA14 with *vfr (PA14_08370)* deleted. Made by mating pLD4042 into LD0. | This study |
| LD732 | PA14 Δ*pqsA-C* | PA14 with *pqsA-C (PA14_51430*, *PA14_51420*, and *PA14_51410)* deleted. | [2] |
| LD1380 | PA14 Δ*rhlR* | PA14 with *rhlR (PA14_19120)* deleted. Made by mating pLD1355 into LD0. | [2] |
| LD4555 | PA14 Δ*antABC* | PA14 with *antABC (PA14_32160*, *PA14_32150*, and *PA14_32140) deleted*. Made by mating pLD4537 into LD0. | This study |
| LD24 | PA14 Δ*phz* (also referred to as Δ*phz1/2*) | PA14 with *phz (PA14_09480-PA14_09410* and *PA14_39970-PA14_39880)* operons deleted. | [3] |
| LD3679 | PA14 Δ*gacA* | PA14 with *gacA (PA14_30650)* deleted. Made by mating pLD3079 into LD0. | This study |
| LD4213 | PA14 Δ*lasR* | PA14 with *lasR (PA14_45960)* deleted. Made by mating pLD4228 into LD0. | This study |
| LD3596 | PA14 Δ*aer1* | PA14 with *aer1 (PA14_44300)* deleted. Made by mating pLD3594 into LD0. | This study |
| LD3604 | PA14 Δ*aer2* | PA14 with *aer2 (PA14_02220)* deleted. Made by mating pLD3579 into LD0. | This study |
| LD3607 | PA14 Δ*aer1* Δ*aer2* | PA14 with *aer1* and *aer2 (PA14_44300* and *PA14_02220)* deleted. Made by mating pLD3594 into LD3604. | This study |
| LD2271 | PA14 Δ*nosP* | PA14 with *nosP (PA14_38990 and PA14_38970)* deleted. Made by mating LD2271 into LD0. | This study |
| LD915 | PA14 Δ*anr* | PA14 with *anr* (*PA14_44490)* deleted. | [4] |
| LD369 | PA14 Δ*pilB* | PA14 with *pilB (PA14_58750)* deleted. | [5] |
| LD1879 | PA14 Δ*pilY1* | PA14 with *pilY1 (PA14_60310)* deleted. Made by mating pLD1858 into LD0. | This study |

| | | | |
|---|---|---|---|
| LD3986 | PA14 Δ*pilA* | PA14 with *pilA (PA14_58730)* deleted. Made by mating pLD3979 into LD0. | This study |
| LD4164 | PA14 Δ*pilT* Δ*pilU* | PA14 with *pilT pilU (PA14_05180 and PA14_05190)* deleted. Made by mating pLD4137 into LD0. | This study |
| LD4644 | PA14 Δ*ssg* | PA14 with *ssg (PA14_66120)* deleted. Made by mating pLD4630 into LD0. | This study |
| LD4913 | PA14 Δ*wapR* | PA14 with *wapR (PA14_66110)* deleted. Made by mating pLD4832 into LD0. | This study |
| LD4635 | PA14 Δ*wbpM* | PA14 with *wbpM (PA14_23470)* deleted. Made by mating pLD4631 into LD0. | This study |
| LD3950 | PA14 Δ*cheY* | PA14 with *cheY (PA14_45620)* deleted. Made by mating pLD3939 into LD0. | This study |
| LD371 | PA14 Δ*flgK* | PA14 with *flgK (PA14_50360)* deleted. | [5] |
| LD384 | PA14 Δ*pilb* Δ*flgK* | PA14 with *pilb* and *flgK (PA14_58750* and *PA14_5036*0) deleted. | [5] |
| LD3621 | PA14 Δ*motA* Δ*motB* | PA14 with *motA* and *motB* (*PA14_65450* and *PA14_65430)* deleted. Made by mating pLD3629 into LD0. | This study |
| LD3622 | PA14 Δ*motC* Δ*motD* | PA14 with *motC and motD (PA14_45560* and *PA14_45540)* deleted. Made by mating pLD3630 into LD0. | This study |
| LD1529 | PA14 Δ*cupA2* | PA14 with *cupA2 (PA14_37040)* deleted. Made by mating pLD into LD0. | This study |
| LD1726 | PA14 ΔcupA Δ*cupD* | PA14 with *cupD1 (PA14_59710)* deleted. Made by mating pLD into LD0. | This study |
| LD3070 | PA14 Δ*pelA-G* | PA14 with *pelA-G (PA14_24480*, *PA14_24490*, *PA14_24500*, *PA14_24510*, *PA14_24530*, *PA14_24550*, and *PA14_24560)* deleted. Made by mating pLDLD3059 into LD0. | This study |
| LD3192 | PA14 Δ*rpoS* | PA14 with *rpoS (PA14_17480)* deleted. Made by mating pLD3471 into LD0. | [6] |
| LD3190 | PA14 Δ*rpoN* | PA14 with *rpoN (PA14_57940)* deleted. Made by mating pLD3473 into LD0. | [6] |
| LD3949 | PA14 Δ*fliA* | PA14 with *fliA (PA14_45630)* deleted. Made by mating pLD3938 into LD0. | This study |

| | | | |
|---|---|---|---|
| LD3674 | PA14 Δ*crc* | PA14 with *crc (PA14_70390)* deleted. | [6] |
| LD3130 | PA14 Δ*ptsP* | PA14 with *ptsP (PA14_04410)* deleted. Made by mating pLD3125 into LD0. | This study |
| LD3694 | PA14 Δ*ptsO* | PA14 with *ptsO* (*PA14_57980)* deleted. Made by mating pLD3635 into LD0. | This study |
| LD3696 | PA14 Δ*ptsO* Δ*ptsP* | PA14 with *ptsO* and *ptsP (PA14_04410* and *PA14_57980)* deleted. Made by mating pLD3635 into LD3130. | This study |
| LD1888 | PA14 Δ*ccoN1* Δ*ccoN2* | PA14 with *ccoN1* and *ccoN2 (PA14_44370* and *PA14_44340)* deleted. Made by mating pLD1610 into LD1784. | [7] |
| LD1976 | PA14 Δ*ccoN1* Δ*ccoN2* Δ*ccoN4* | PA14 with *ccoN1*, *ccoN2*, and *ccoN4 (PA14_443470*, *PA14_44340*, and *PA14_10500)* deleted. Made by mating pLD1264 into LD1888. | [7] |
| LD1933 | PA14 Δ*cco1* Δ*cco2* | PA14 with both *cco* operons *(PA14_44340-PA14_44400*) deleted simultaneously. | [7] |
| LD3196 | PA14 Δ*dipA* | PA14 with *dipA (PA14_66320)* deleted. Made by mating pLD1204 into LD0. | This study |
| LD4687 | PA14 Δ*relA* | PA14 with *relA (PA14_52180)* deleted. Made by mating pLD4642 into LD0. | [8] |
| LD2177 | PA14 Δ*sadC* | PA14 with *sadC (PA14_56280)* deleted. Made by mating pLD2173 into LD0. | [8] |
| LD2569 | PA14 Δ*bifA* | PA14 with *bifA (PA14_56790)* deleted. Made by mating pLD2565 into LD0. | [8] |
| LD2183 | PA14 Δ*roeA* | PA14 with *roeA (PA14_50060)* deleted. Made by mating pLD2179 into LD0. | [8] |
| LD2227 | PA14 Δ*rmcA* | PA14 with *rmcA* (*PA14_07500)* deleted. Made by mating pLD909 into LD0. | [8] |
| LD2428 | PA14 Δ*wspR* | PA14 with *wspR* (*PA14_16500)* deleted. Made by mating pLD4040 into LD0. | [8] |
| LD3917 | PA14 Δ*cyaA* | PA14 with *cyaA (PA14_69610)* deleted. Made by mating pLD3910 into LD0. | This study |
| LD3920 | PA14 Δ*cyaB* | PA14 with *cyaB (PA14_22620)* deleted. Made by mating pLD3911 into LD0. | This study |

| | | | |
|---|---|---|---|
| LD3923 | PA14 Δ*cyaA* Δ*cyaB* | PA14 with *cyaA cyaB (PA14_69610* and *PA14_22620)* deleted. Made by mating pLD3910 into LD3920 LD0. | This study |
| LD3914 | PA14 Δ*cpdA* | PA14 with *cpdA (PA14_65690)* deleted. Made by mating pLD3909 into LD0. | This study |
| LD3625 | PA14 Δ*ackA* | PA14 with *ackA (PA14_53470)* deleted. Made by mating pLD3609 into LD0. | This study |
| LD2729 | PA14 Δ*ldhA* | PA14 with *ldhA (PA14_52270)* deleted. Made by mating pLD2728 into LD0. | [9] |
| LD3646 | PA14 Δ*ldhA* Δ*ackA* | PA14 with *ldhA ackA (PA14_53470* and *PA14_52270)* deleted. Made by mating pLD2728 into LD3625. | This study |
| LD4833 | PA14 Δ*pilA* Δ*ssg* | PA14 with *pilA ssg (PA14_58730* and *PA14_66120)* deleted. Made by mating pLD4630 into LD3986. | This study |
| LD4836 | PA14 Δ*pilA* Δ*wbpM* | PA14 with *pilA wbpM (PA14_58730* and *PA14_23470)* deleted. Made by mating pLD4631 into LD3986. | This study |
| LD4839 | PA14 Δ*ssg* Δ*wbpM* | PA14 with *ssg wbpM (PA14_66120* and *PA14_23470)* deleted. Made by mating pLD4631 into LD4644. | This study |
| LD4835 | PA14 Δ*wbpM* Δ*ssg* | PA14 with *wbpM ssg (PA14_23470* and *PA14_66120)* deleted. Made by mating pLD4630 into LD4635. | This study |
| LD4764 | PA14 $P_{PA1/04/03}$-mScarlet | PA14 constitutively expressing mScarlet. Made by mating pLD3433 into LD0. | This study |
| LD4070 | PA14 Δ*vfr* $P_{PA1/04/03}$-mScarlet | PA14 Δ*vfr* constitutively expressing mScarlet. Made by mating pLD3433 into LD4058. | This study |
| LD4007 | PA14 Δ*pqsA-C* $P_{PA1/04/03}$-mScarlet | PA14 Δ*pqsA-C* constitutively expressing mScarlet. Made by mating pLD3433 into LD732. | This study |
| LD4008 | PA14 Δ*rhlR* $P_{PA1/04/03}$-mScarlet | PA14 Δ*rhlR* constitutively expressing mScarlet. Made by mating pLD3433 into LD1380. | This study |
| LD4765 | PA14 Δ*antABC* $P_{PA1/04/03}$-mScarlet | PA14 Δ*antABC* constitutively expressing mScarlet. Made by mating pLD3433 into LD4555. | This study |
| LD3608 | PA14 Δ*phz* (also | PA14Δ*phz* constitutively expressing mScarlet. | This study |

| | | | |
|---|---|---|---|
| | referred to as Δ*phz1/2*) $P_{PA1/04/03}$-mScarlet | Made by mating pLD3433 into LD24. | |
| LD5074 | PA14 Δ*gacA* $P_{PA1/04/03}$-mScarlet | PA14 Δ*gacA* constitutively expressing mScarlet. Made by mating pLD3433 into LD3679. | This study |
| LD4763 | PA14 Δ*lasR* $P_{PA1/04/03}$-mScarlet | PA14 Δ*lasR* constitutively expressing mScarlet. Made by mating pLD3433 into LD4213. | This study |
| LD5075 | PA14 Δ*aer1* $P_{PA1/04/03}$-mScarlet | PA14 Δ*aer1* constitutively expressing mScarlet. Made by mating pLD3433 into LD3596. | This study |
| LD5076 | PA14 Δ*aer2* $P_{PA1/04/03}$-mScarlet | PA14 Δ*aer2* constitutively expressing mScarlet. Made by mating pLD3433 into LD3604. | This study |
| LD5077 | PA14 Δ*aer1* Δ*aer2* $P_{PA1/04/03}$-mScarlet | PA14 Δ*aer1* Δ*aer2* constitutively expressing mScarlet. Made by mating pLD3433 into LD3607. | This study |
| LD4217 | PA14 Δ*nosP* $P_{PA1/04/03}$-mScarlet | PA14 Δ*nosP* constitutively expressing mScarlet. Made by mating pLD3433 into LD. | This study |
| LD4127 | PA14 Δ*anr* $P_{PA1/04/03}$-mScarlet | PA14 Δ*anr* constitutively expressing mScarlet. Made by mating pLD3433 into LD915. | This study |
| LD5078 | PA14 Δ*pilB* $P_{PA1/04/03}$-mScarlet | PA14 Δ*pilB* constitutively expressing mScarlet. Made by mating pLD3433 into LD369. | This study |
| LD5079 | PA14 Δ*pilY1* $P_{PA1/04/03}$-mScarlet | PA14 Δ*pilY1* constitutively expressing mScarlet. Made by mating pLD3433 into LD1879. | This study |
| LD3987 | PA14 Δ*pilA* $P_{PA1/04/03}$-mScarlet | PA14 Δ*pilA* constitutively expressing mScarlet. Made by mating pLD3433 into LD3986. | This study |
| LD4175 | PA14 Δ*pilT* Δ*pilU* $P_{PA1/04/03}$-mScarlet | PA14 Δ*pilT* Δ*pilU* constitutively expressing mScarlet. Made by mating pLD3433 into LD4164. | This study |
| LD4665 | PA14 Δ*ssg* $P_{PA1/04/03}$-mScarlet | PA14 Δ*ssg* constitutively expressing mScarlet. Made by mating pLD3433 into LD4644. | This study |
| LD4921 | PA14 Δ*wapR* $P_{PA1/04/03}$-mScarlet | PA14 Δ*wapR* constitutively expressing mScarlet. Made by mating pLD3433 into LD4913. | This study |

| | | | |
|---|---|---|---|
| LD4666 | PA14 Δ*wbpM* $P_{PA1/04/03}$-mScarlet | PA14 Δ*wbpM* constitutively expressing mScarlet. Made by mating pLD3433 into LD4635. | This study |
| LD3953 | PA14 Δ*cheY* $P_{PA1/04/03}$-mScarlet | PA14 Δ*cheY* constitutively expressing mScarlet. Made by mating pLD3433 into LD3950. | This study |
| LD5080 | PA14 Δ*flgK* $P_{PA1/04/03}$-mScarlet | PA14 Δ*flgK* constitutively expressing mScarlet. Made by mating pLD3433 into LD371. | This study |
| LD5081 | PA14 Δ*pilb* Δ*flgK* $P_{PA1/04/03}$-mScarlet | PA14 Δ*pilb* Δ*flgK* constitutively expressing mScarlet. Made by mating pLD3433 into LD384. | This study |
| LD5082 | PA14 Δ*motA* Δ*motB* $P_{PA1/04/03}$-mScarlet | PA14 Δ*motA* Δ*motB* constitutively expressing mScarlet. Made by mating pLD3433 into LD3621. | This study |
| LD5083 | PA14 Δ*motC* Δ*motD* $P_{PA1/04/03}$-mScarlet | PA14 Δ*motC* Δ*motD* constitutively expressing mScarlet. Made by mating pLD3433 into LD3622. | This study |
| LD5084 | PA14 Δ*cupA2* $P_{PA1/04/03}$-mScarlet | PA14 Δ*cupA2* constitutively expressing mScarlet. Made by mating pLD3433 into LD1529. | This study |
| LD5085 | PA14 Δ*cupA* Δ*cupD* $P_{PA1/04/03}$-mScarlet | PA14 Δ*cupA* Δ*cupD* constitutively expressing mScarlet. Made by mating pLD3433 into LD1726. | This study |
| LD5086 | PA14 Δ*pelA-G* $P_{PA1/04/03}$-mScarlet | PA14 Δ*pelA-G* constitutively expressing mScarlet. Made by mating pLD3433 into LD3070. | This study |
| LD5087 | PA14 Δ*rpoS* $P_{PA1/04/03}$-mScarlet | PA14 Δ*rpoS* constitutively expressing mScarlet. Made by mating pLD3433 into LD3192. | This study |
| LD5088 | PA14 Δ*rpoN* $P_{PA1/04/03}$-mScarlet | PA14 Δ*rpoN* constitutively expressing mScarlet. Made by mating pLD3433 into LD3190. | This study |
| LD3951 | PA14 Δ*fliA* $P_{PA1/04/03}$-mScarlet | PA14 Δ*fliA* constitutively expressing mScarlet. Made by mating pLD3433 into LD3949. | This study |
| LD5089 | PA14 Δ*crc* $P_{PA1/04/03}$-mScarlet | PA14 Δ*crc* constitutively expressing mScarlet. Made by mating pLD3433 into LD3674. | This study |
| LD5090 | PA14 Δ*ptsP* | PA14 Δ*ptsP* constitutively expressing mScarlet. | This study |

| | | | |
|---|---|---|---|
| | $P_{PA1/04/03}$-mScarlet | Made by mating pLD3433 into LD3130. | |
| LD4008 | PA14 Δ*ptsO* $P_{PA1/04/03}$-mScarlet | PA14 Δ*ptsO* constitutively expressing mScarlet. Made by mating pLD3433 into LD3694. | This study |
| LD4009 | PA14 Δ*ptsO* Δ*ptsP* $P_{PA1/04/03}$-mScarlet | PA14 Δ*ptsO* Δ*ptsP* constitutively expressing mScarlet. Made by mating pLD3433 into LD3696. | This study |
| LD2013 | PA14 Δ*ccoN1* Δ*ccoN2* $P_{PA1/04/03}$-YFP | PA14 Δ*ccoN1* Δ*ccoN2* constitutively expressing YFP. Made by mating pLD3433 into LD1888. | [7] |
| LD2136 | PA14 Δ*ccoN1* Δ*ccoN2* Δ*ccoN4* $P_{PA1/04/03}$-YFP | PA14 Δ*ccoN1* Δ*ccoN2* Δ*ccoN4* constitutively expressing YFP. Made by mating pLD3433 into LD1976. | [7] |
| LD2012 | PA14 Δ*cco1* Δ*cco2* $P_{PA1/04/03}$-YFP | PA14 Δ*cco1* Δ*cco* constitutively expressing YFP. Made by mating pLD3433 into LD1933. | [7] |
| LD5091 | PA14 Δ*dipA* $P_{PA1/04/03}$-mScarlet | PA14 Δ*dipA* constitutively expressing mScarlet. Made by mating pLD3433 into LD3196. | This study |
| LD4766 | PA14 Δ*relA* $P_{PA1/04/03}$-mScarlet | PA14 Δ*relA* constitutively expressing mScarlet. Made by mating pLD3433 into LD4687. | This study |
| LD2925 | PA14 Δ*sadC* $P_{PA1/04/03}$-YFP | PA14 Δ*sadC* constitutively expressing YFP. Made by mating pLD3433 into LD2177. | This study |
| LD2773 | PA14 Δ*bifA* $P_{PA1/04/03}$-YFP | PA14 Δ*bifA* constitutively expressing YFP. Made by mating pLD3433 into LD2569. | This study |
| LD2183 | PA14 Δ*roeA* $P_{PA1/04/03}$-YFP | PA14 Δ*roeA* constitutively expressing YFP. Made by mating pLD3433 into LD2183. | This study |
| LD2775 | PA14 Δ*rmcA* $P_{PA1/04/03}$-YFP | PA14 Δ*rmcA* constitutively expressing YFP. Made by mating pLD3433 into LD2227. | This study |
| LD2488 | PA14 Δ*wspR* $P_{PA1/04/03}$-YFP | PA14 Δ*wspR* constitutively expressing YFP. Made by mating pLD3433 into LD2428. | This study |
| LD4069 | PA14 Δ*cyaA* Δ*cyaB* $P_{PA1/04/03}$-mScarlet | PA14 Δ*cyaA* Δ*cyaB* constitutively expressing mScarlet. Made by mating pLD3433 into LD3923. | This study |
| LD4062 | PA14 Δ*cpdA* $P_{PA1/04/03}$-mScarlet | PA14 Δ*cpdA* constitutively expressing mScarlet. Made by mating pLD3433 into LD3914. | This study |

| | | | |
|---|---|---|---|
| LD5092 | PA14 Δ*ackA* $P_{PA1/04/03}$-mScarlet | PA14 Δ*ackA* constitutively expressing mScarlet. Made by mating pLD3433 into LD3625. | This study |
| LD5093 | PA14 Δ*ldhA* $P_{PA1/04/03}$-mScarlet | PA14 Δ*ldhA* constitutively expressing mScarlet. Made by mating pLD3433 into LD2729. | This study |
| LD5094 | PA14 Δ*pilA* Δ*ssg* $P_{PA1/04/03}$-mScarlet | PA14 Δ*pilA* Δ*ssg* constitutively expressing mScarlet. Made by mating pLD3433 into LD3646. | This study |
| LD4850 | PA14 Δ*pilA* Δ*wbpM* $P_{PA1/04/03}$-mScarlet | PA14 Δ*pilA* Δ*wbpM* constitutively expressing mScarlet. Made by mating pLD3433 into LD4833. | This study |
| LD4853 | PA14 Δ*ldhA* Δ*ackA* $P_{PA1/04/03}$-mScarlet | PA14 Δ*ldhA* Δ*ackA* constitutively expressing mScarlet. Made by mating pLD3433 into LD4836. | This study |
| LD4852 | PA14 Δ*wbpM* Δ*ssg* $P_{PA1/04/03}$-mScarlet | PA14 Δ*wbpM* Δ*ssg* constitutively expressing mScarlet. Made by mating pLD3433 into LD4835. | This study |
| LD4855 | PA14 Δ*ssg* Δ*wbpM* $P_{PA1/04/03}$-mScarlet | PA14 Δ*ssg* Δ*wbpM* constitutively expressing mScarlet. Made by mating pLD3433 into LD4839. | This study |
| LD3801 | WT $P_{PA1/04/03}$-eGFP | PA14 constitutively expressing eGFP. Made by mating pLD3655 into LD0. | This study |
| LD4292 | Δ*pilA* $P_{PA1/04/03}$-eGFP | PA14 constitutively expressing eGFP. Made by mating pLD3655 into LD3986. | This study |
| LD4842 | Δ*wbpM* $P_{PA1/04/03}$-eGFP | PA14 constitutively expressing eGFP. Made by mating pLD3655 into LD4635. | This study |
| 5047 | Δ*wbpM* Δ*pilA* $P_{PA1/04/03}$-eGFP | PA14 constitutively expressing eGFP. Made by mating pLD3655 into LD4836. | This study |
| LD4291 | Δ*gacA* $P_{PA1/04/03}$-eGFP | PA14 constitutively expressing eGFP. Made by mating pLD3655 into LD3679. | This study |
| LD4373 | WT attB::rhaSR-PrhaBAD-mScarlet | attB::rhaSR-PrhaBAD-mScarlet. Made by mating pLD4358 into LD0. | This study |
| LD4374 | Δ*gacA* attB::rhaSR-PrhaBAD-mScarlet | attB::rhaSR-PrhaBAD-mScarlet. Made by mating pLD4358 into LD3679. | This study |

| | | | |
|---|---|---|---|
| LD4376 | ∆*pilA* attB::rhaSR-PrhaBAD-mScarlet | attB::rhaSR-PrhaBAD-mScarlet. Made by mating pLD4358 into LD3986. | This study |
| LD4992 | ∆*wbpM* attB::rhaSR-PrhaBAD-mScarlet | attB::rhaSR-PrhaBAD-mScarlet. Made by mating pLD4358 into LD4635. | This study |
| LD5052 | ∆*wbpM* ∆*pilA* attB::rhaSR-PrhaBAD-mScarlet | attB::rhaSR-PrhaBAD-mScarlet. Made by mating pLD4358 into LD4836. | This study |
| *E. coli* strains | | | |
| LD44 | UQ950 | E. coli DH5α λ(pir) host for cloning; F-Δ(*argF-lac*)169 Φ80 d*lacZ58*(ΔM15) *glnV44*(AS) *rfbD1 gyrA96*(NalR) *recA1 endA1 spoT1 thi-1 hsdR17 deoR* λpir+ | D. Lies |
| LD661 | BW29427 | Donor strain for conjugation: *thrB1004 pro thi rpsL hsdS lacZ* ΔM15RP4–1360 Δ(*araBAD*)567 Δ*dapA1341*::[*erm pir*(wt)] | W. Metcalf |
| LD69 | β2155 | Helper strain. *thrB1004 pro thi strA hsdsS lacZ*∆M15 (*F' lacZ∆M15 lacI*$^q$ *tra∆36 proA*$^+$ *proB*$^+$) ∆*dapA::erm* (Erm$^r$)*pir::*RP4 [*::kan* (Km$^r$) from SM10] | [10] |
| LD2901 | S17-1 | Str$^R$, Tp$^R$, F$^-$ RP4-2-Tc::Mu *aphA*::Tn7 *recA* λpir lysogen | R. Simon |
| *Saccharomyces cerevisiae* strains | | | |
| LD676 | InvSc1 | *MATα/MATα leu2/leu2 trp1-289/trp1-289 ura3-52/ura3-52 his3-Δ1/his3-Δ1* | Invitrogen |

**Table S2. Plasmids used in this study.**

| Plasmid Name | Description | Source |
|---|---|---|
| pMQ30 | Yeast-based allelic-exchange vector; *sacB*$^+$, CEN/ARSH, URA3$^+$, Gm$^R$. | [11] |
| pFLP2 | Site-specific excision vector with cI857-controlled FLP recombinase. encoding sequence, sacB$^+$, Amp$^R$. Used to insert LD2722-based plasmids into P. aeruginosa strains. | [12] |

| | | |
|---|---|---|
| pLD2722 | Gm$^R$, Tet$^R$ flanked by Flp recombinase target (FRT) sites to resolve out resistance cassettes. | [7] |
| pAKN69-YFP | Gm$^R$, Cm$^R$ mini-Tn7 $P_{PA1/04/03}$*::yfp* | [13] |
| pAKN69-mScarlet | Gm$^R$, Cm$^R$ mini-Tn7 $P_{PA1/04/03}$*::mScarlet* | This study |
| pLD4042 | Δ*vfr (PA14_08370)* PCR fragment introduced into pMQ30 by gap repair cloning in yeast strain InvSc1. | This study |
| pLD4537 | Δ*antABC (PA14_32160*, *PA14_32150*, and *PA14_32140)* PCR fragment introduced into pMQ30 by gap repair cloning in yeast strain InvSc1. | This study |
| pLD3079 | Δ*gacA (PA14_30650)* PCR fragment introduced into pMQ30 by gap repair cloning in yeast strain InvSc1. | This study |
| pLD4228 | Δ*lasR (PA14_45960)* PCR fragment introduced into pMQ30 by gap repair cloning in yeast strain InvSc1. | This study |
| pLD3594 | Δ*aer1 (PA14_44300)* PCR fragment introduced into pMQ30 by gap repair cloning in yeast strain InvSc1. | This study |
| pLD3579 | Δ*aer2 (PA14_02220)* PCR fragment introduced into pMQ30 by gap repair cloning in yeast strain InvSc1. | This study |
| pLD3594 | Δ*aer1* Δ*aer2 (PA14_44300* and *PA14_02220)* PCR fragment introduced into pMQ30 by gap repair cloning in yeast strain InvSc1. | This study |
| pLD3979 | Δ*pilA (PA14_58730)* PCR fragment introduced into pMQ30 by gap repair cloning in yeast strain InvSc1. | This study |
| pLD4137 | Δ*pilT* Δ*pilU (PA14_05180 and PA14_05190)* PCR fragment introduced into pMQ30 by gap repair cloning in yeast strain InvSc1. | This study |
| pLD4630 | Δ*ssg (PA14_66120)* PCR fragment introduced into pMQ30 by gap repair cloning in yeast strain InvSc1. | This study |
| pLD4832 | Δ*wapR (PA14_66110)* PCR fragment introduced into pMQ30 by gap repair cloning in yeast strain InvSc1. | This study |
| pLD4631 | Δ*wbpM (PA14_23470)* PCR fragment introduced into pMQ30 by gap repair cloning in yeast strain InvSc1. | This study |
| pLD3939 | Δ*cheY (PA14_45620)* PCR fragment introduced into pMQ30 by gap repair cloning in yeast strain InvSc1. | This study |
| pLD3629 | Δ*motA* Δ*motB* (*PA14_65450* and *PA14_65430)* PCR fragment introduced into pMQ30 by gap repair cloning in yeast strain | This study |

| | | |
|---|---|---|
| | InvSc1. | |
| pLD3630 | ∆*motC* ∆*motD (PA14_45560* and *PA14_45540)* PCR fragment introduced into pMQ30 by gap repair cloning in yeast strain InvSc1. | This study |
| pLD3938 | ∆*fliA (PA14_45630)* PCR fragment introduced into pMQ30 by gap repair cloning in yeast strain InvSc1. | This study |
| pLD3635 | ∆*ptsP (PA14_04410)* PCR fragment introduced into pMQ30 by gap repair cloning in yeast strain InvSc1. | This study |
| pLD1204 | ∆*dipA (PA14_66320)* PCR fragment introduced into pMQ30 by gap repair cloning in yeast strain InvSc1. | This study |
| pLD3910 | ∆*cyaA PA14_69610* PCR fragment introduced into pMQ30 by gap repair cloning in yeast strain InvSc1. | This study |
| pLD3909 | ∆*cpdA (PA14_65690)* PCR fragment introduced into pMQ30 by gap repair cloning in yeast strain InvSc1. | This study |
| pLD3609 | ∆*ackA (PA14_53470)* PCR fragment introduced into pMQ30 by gap repair cloning in yeast strain InvSc1. | This study |
| pLD4358 | pSEK109-rhaSR-PrhaBAD | This study |

**Table S3. Primers used in this study.**

| Primer Number | Sequence |
|---|---|
| Primers for plasmid pLD4042 (used to make ∆*vfr*) | |
| 3611 | cctgcaggtcgactctagagGGGAGAAGATGGACGAACTG |
| 3612 | CTACACCGCAAAGAGCACCCTGGTGCATGTGAAAGGAAA |
| 3613 | TTTCCTTTCACATGCACCAGGGTGCTCTTTGCGGTGTAG |
| 3614 | cagctatgaccatgattacg GATGGCGATTGTGGTTTGTT |
| Primers for plasmid pLD4537 (used to make ∆*antABC*) | |
| 3962 | ggcaaattctgttttatcagaccgcttctgcgttctgatTCAGTCCCTTGATCAGCGC |
| 3963 | GTCGAAATGCTCGTGCAGGTACAGCTCCGGTTCGGTGAAC |
| 3964 | GTTCACCGAACCGGAGCTGTACCTGCACGAGCATTTCGAC |
| 3965 | aattgtgagcggataacaatttcacacaggaaacagctCCCGGCGAATTTCTACTACGA |

| | |
|---|---|
| Primers for plasmid pLD3079 (used to make Δ*gacA*) | |
| 1932 | ccaggcaaattctgttttatcagaccgcttctgcgttctgatAGCGTGCGACGTTATGTCT |
| 1933 | gaagatgcggtagcgatagACCACTTGCAAGCCTTCG |
| 1934 | cgaaggcttgcaagtggtCTATCGCTACCGCATCTTC |
| 1935 | ggaattgtgagcggataacaatttcacacaggaaacagctTCACGTTGACGATCAACTCC |
| Primers for plasmid pLD4228 (used to make Δ*lasR*) | |
| 3813 | cctgcaggtcgactctagagACAGGTCCCCGTCATGAAAC |
| 3814 | GTTTTCTTGAGCTGGAACGCATGGCCGTTAATTTGGGTCT |
| 3815 | AGACCCAAATTAACGGCCATGCGTTCCAGCTCAAGAAAAC |
| 3816 | cagctatgaccatgattacgGATCAACATGGTCACCTCCA |
| Primers for plasmid pLD3594 (used to make Δ*aer1*) | |
| 2958 | aggcaaattctgttttatcagaccgcttctgcgttctgatTGGAACCGAAGGACTTCAAC |
| 2942 | CTCGTCGGAGGTCTGCTCTTTGAGATTGGTGGTGGTGA |
| 2943 | TCACCACCACCAATCTCAAAGAGCAGACCTCCGACGAG |
| 2944 | ggaattgtgagcggataacaatttcacacaggaaacagctAGATCCTCTGCGGCGATT |
| Primers for plasmid pLD3579 (used to make Δ*aer2*) | |
| 2950 | aggcaaattctgttttatcagaccgcttctgcgttctgatCACGGTGACGATCACCAGGT |
| 2951 | CTCCAGTCGTTCGCCGACAGCGTGGTCCAGCTCGCC |
| 2952 | GGCGAGCTGGACCACGCTGTCGGCGAACGACTGGAG |
| 2953 | ggaattgtgagcggataacaatttcacacaggaaacagctCTGGCAACTGGTGTTCATGC |
| Primers for plasmid pLD3979 (used to make Δ*pilA*) | |
| 3583 | aggcaaattctgttttatcagaccgcttctgcgttctgatCCTCACCCTCTGAACGAATC |
| 3584 | GATTACACCATTCGTGCTCG ATCCAATGCGGTTTCCAGT |
| 3585 | ACTGGAAACCGCATTGGAT CGAGCACGAATGGTGTAATC |
| 3586 | ggaattgtgagcggataacaatttcacacaggaaacagctACTTCATGGAGCATGCCATC |
| Primers for plasmid pLD4137 (used to make Δ*pilT* Δ*pilU*) | |
| 3666 | aggcaaattctgttttatcagaccgcttctgcgttctgatGAACGCTATGCGGGGTGT |
| 3667 | GTCAGGTTCGACAGGTGGTC GGAGAGGTGCAGGTCCGAAG |
| 3668 | CTTCGGACCTGCACCTCTCC GACCACCTGTCGAACCTGAC |
| 3669 | ggaattgtgagcggataacaatttcacacaggaaacagctGCCTATCCTGTAGGCTGGTG |
| Primers for plasmid pLD4630 (used to make Δ*ssg*) | |

| | |
|---|---|
| 4193 | aggcaaattctgttttatcagaccgcttctgcgttctgatAGAAAAACAACCGCGCAAT |
| 4194 | GGCTACTTCCGCAAGCACGCCGCTGTCGGTCAGAGTATC |
| 4195 | GATACTCTGACCGACAGCGGCGTGCTTGCGGAAGTAGCC |
| 4196 | ggaattgtgagcggataacaatttcacacaggaaacagctTACCTGAACCTGACGGGACT |
| Primers for plasmid pLD4832 (used to make ∆*wapR*) | |
| 4349 | aggcaaattctgttttatcagaccgcttctgcgttctgatGGCATGATCGTGGCTTATATC |
| 4350 | CAGACGTACCGCAACGTCTCGAACCGGGACAAGGT |
| 4351 | ACCTTGTCCCGGTTCGAGACGTTGCGGTACGTCTG |
| 4352 | ggaattgtgagcggataacaatttcacacaggaaacagctAAGAATTTCGAGGCGATGG |
| Primers for plasmid pLD4631 (used to make ∆*wbpM* ) | |
| 4201 | aggcaaattctgttttatcagaccgcttctgcgttctgatCATGGATGGCATTGATGGTA |
| 4202 | GTAGTCGTCCTTCTCCACGGCGAAGGCTAGCCAGAGCGATA |
| 4203 | TATCGCTCTGGCTAGCCTTCGCCGTGGAGAAGGACGACTAC |
| 4204 | ggaattgtgagcggataacaatttcacacaggaaacagctGGTGATCCGCACCAACTATT |
| Primers for plasmid pLD3939 (used to make ∆*cheY*) | |
| 3424 | aggcaaattctgttttatcagaccgcttctgcgttctgat CCCAGAGACGTGTTACTCGAA |
| 3425 | TCAACGGCTATGTGGTCAAAGGACTTGGGCTTCACCAATA |
| 3426 | TATTGGTGAAGCCCAAGTCCTCAACGGCTATGTGGTCAAA |
| 3427 | ggaattgtgagcggataacaatttcacacaggaaacagctGAGCCGTATACGAGGCATTC |
| Primers for plasmid pLD3629 (used to make ∆*motA* ∆*motB*) | |
| 2994 | aggcaaattctgttttatcagaccgcttctgcgttctgatGCATCGAAAAGATCCAGGAA |
| 2995 | ATCCAGCCCTTCGAGGTCGTGCAGAAGAAGCTCAACC |
| 2996 | GGTTGAGCTTCTTCTGCACGACCTCGAAGGGCTGGAT |
| 2997 | ggaattgtgagcggataacaatttcacacaggaaacagctACCTGCTGCTGGACTTCG |
| Primers for plasmid pLD3630 (used to make ∆*motC* ∆*motD*) | |
| 3002 | aggcaaattctgttttatcagaccgcttctgcgttctgatGTAGGACTGCAAGGCCAGTT |
| 3003 | ATCATCCTGGCGTTCGTCGGCACGCAAGCTGCAGATACT |
| 3004 | AGTATCTGCAGCTTGCGTGCCGACGAACGCCAGGATGAT |
| 3005 | ggaattgtgagcggataacaatttcacacaggaaacagctCCGGTCCTGATGTTCTCCT |
| Primers for plasmid pLD3938 (used to make ∆*fliA*) | |
| 3484 | aggcaaattctgttttatcagaccgcttctgcgttctgatGAACTGTTCGATGCGCTTG |

| | |
|---|---|
| 3485 | CATCGCCTATCACCTGCTCGAGGAGATCGGCGAGGTACTG |
| 3486 | CAGTACCTCGCCGATCTCCTCGAGCAGGTGATAGGCGATG |
| 3487 | ggaattgtgagcggataacaatttcacacaggaaacagctCATCGTTCCCGCCAATAC |
| Primers for plasmid pLD3635 (used to make Δ*ptsP*) | |
| 2475 | aggcaaattctgttttatcagaccgcttctgcgttctgatCGGCTCGATCACTTCCTGCA |
| 2476 | ctgcgggttgtcgaaggtgagCATGGCTTCCTTGACCCGCTG |
| 2477 | cagcgggtcaaggaagccatgCTCACCTTCGACAACCCGCAG |
| 2478 | ggaattgtgagcggataacaatttcacacaggaaacagctGATGAACTCCTCGCCGCCC |
| Primers for plasmid pLD1204 (used to make Δ*dipA*) | |
| 2611 | ggaattgtgagcggataacaatttcacacaggaaacagctGCAGCAACTGGTGAAGCGC |
| 2612 | GCAGCTCGCTGAGGTTCTcgcGCGCCGGAAGCTCGGA |
| 2613 | TCCGAGCTTCCGGCGCgcgAGAACCTCAGCGAGCTGCT |
| 2614 | ccaggcaaattctgttttatcagaccgcttctgcgttctgatATCTGCAGCTCGCCCAG |
| Primers for plasmid pLD3910 (used to make Δ*cyaA*) | |
| 3502 | cctgcaggtcgactctagagCAGCTCAATCCCTACCCCTA |
| 3503 | TAACGCAGATAGTGCAGCGTGTCGAGATCGAGGCTGAGTG |
| 3504 | CACTCAGCCTCGATCTCGACACGCTGCACTATCTGCGTTA |
| 3505 | cagctatgaccatgattacgAAGGCAAGGTCTCGATCCTC |
| Primers for plasmid pLD3909 (used to make Δ*cpdA*) | |
| 3494 | cctgcaggtcgactctagagGTCGATGACTTCCAGCGAGT |
| 3495 | CGTACTGCTGGTGCAGCTTTTCGAAGTGGACTACGACACC |
| 3496 | GGTGTCGTAGTCCACTTCGAAAAGCTGCACCAGCAGTACG |
| 3497 | cagctatgaccatgattacg CCTGATCGACAAGGACGAG |
| Primers for plasmid pLD3609 (used to make Δ*ackA*) | |
| 3022 | caggcaaattctgttttatcagaccgcttctgcgttctgatCAATCGGCGAATGAACGCCG |
| 3023 | GGATCACCAGTACCCGCGGGTGCAGGGGAAACAGGGAGT |
| 3024 | ACTCCCTGTTTCCCCTGCACCCGCGGGTACTGGTGATCC |
| 3025 | ggaattgtgagcggataacaatttcacacaggaaacagctCGTGCGCTGAGCAGATCG |
| Primers for plasmid pLD4358 (used to make pSEK109-rhaSR-PrhaBAD) | |
| 3920 | gattcgactgcactagtTTAATCTTTCTGCGAATTGAGATGAC |
| 3921 | gattcgactgcctcgagTACGACCAGTCTAAAAAGCGC |
